# Single-Molecule Analysis of DNA Base-Stacking Energetics Using Patterned DNA Nanostructures

**DOI:** 10.1101/2022.09.08.506950

**Authors:** Abhinav Banerjee, Micky Anand, Simanta Kalita, Mahipal Ganji

## Abstract

DNA double helix structure is stabilized by the base-pairing and the base-stacking interactions. Base-stacking interactions originating from hydrophobic interactions between the nucleobases predominantly contribute to the duplex stability. A comprehensive understanding of dinucleotide base-stacking interactions is lacking owing to the unavailability of sensitive techniques that can measure these weak interactions. Earlier studies attempting to address this question only managed to estimate the base-pair stacking interactions, however, disentangling individual base-stacking interactions was enigmatic. By combining multiplexed DNA-PAINT imaging with designer DNA nanostructures, we experimentally measure the free energy of dinucleotide base-stacking at the single-molecule level. Multiplexed imaging enabled us to extract binding kinetics of an imager strand with and without additional dinucleotide stacking interactions in a single imaging experiment, abolishing any effects of experimental variations. The DNA-PAINT data showed that a single additional dinucleotide base-stacking results in as much as 250-fold stabilization of the imager strand binding. We found that the dinucleotide base-stacking energies vary from -1.18 ± 0.17 kcal/mol to -3.57 ± 0.08 kcal/mol for C|T and A|C base-stackings, respectively. We demonstrate the application of base-stacking energetics in designing DNA-PAINT probes for multiplexed super-resolution imaging. Our results will aid in designing functional DNA nanostructures, DNA and RNA aptamers, and facilitate better predictions of the local DNA structure.

## INTRODUCTION

DNA as a genetic material undergoes constant deformations for fulfilling the cellular needs, yet the genetic material is efficiently transferred to many generations without any substantial mutations. This calls for robust local interactions within the molecule to ensure long-term stability. The DNA double helix achieves this thermodynamic stability by the base-pairing between the complementary nucleotides^1^ and base-stacking interactions between the adjacent nucleotides.^2^ Biochemical analysis suggests that base-stacking energies predominantly contribute to the stabilization of DNA as compared to the base-pairing interactions.^3^ The base-stacking interactions play a role in nucleic acids metabolic processes,^2, 4, 5^ as well as aid in designing functional hierarchical DNA nanostructures, and DNA and RNA aptamers.^6-8^,

To date base-stacking energetics were measured from bulk biochemical studies using thermal denaturing of DNA duplexes.^9-17^ Based on these data, a unified nearest neighbor (NN) model has been developed, which predicts the sequence-dependent DNA thermal stability with an accuracy of 2 °C.^18^ However, the NN approximation does not separate base-pairing and base-stacking interactions. Earlier attempts to estimate the base-stacking energies utilized biochemical studies analyzing the relative electrophoresis of DNA molecules having nicks or gaps on a urea polyacrylamide gels.^3, 19-21^ More recently, single-molecule optical tweezer experiments measured the force-dependent dissociation rate between the blunt-ends of the parallel DNA beams to derive individual base-pair stacking energetics.^22^ This assay only estimated free energies by means of extrapolating the force applied across DNA beams consisting of several DNA blunt ends. However, direct measurement of individual base-stacking forces, especially at the single-molecule level, between dinucleotides was not conceivable due to the unavailability of sensitive experimental techniques that can capture these weak interactions.

In this report, we measure individual dinucleotide base-stacking energetics using a Single-Molecule Localization Microscopy (SMLM) technique called DNA-PAINT exploiting DNA nanotechnology for multiplexing.^23-25^ DNA-PAINT enables us to directly access the binding dynamics of a fluorophore-labeled short oligonucleotide (denoted as imager) to a complementary strand (denoted as docking strand) positioned on a DNA origami nanostructure.^25, 26^ We compare the imager’s binding kinetics with stacking and non-stacking configurations at its terminal nucleotide. We utilized patterned DNA origami nanostructures for multiplexed imaging of stacked and non-stacked configurations in a single experiment under equilibrium conditions, thus, avoiding any experimental variations such as temperature and buffer composition. Our kinetic analysis of single-molecule data resulted in an unexpectedly wide range of dwell time stabilizations, varying from 7-to 250-folds for C|T and A|C, respectively, by individual base-stacking interactions. We present the free energy of all 16 possible base-stacking interactions ranging from 1.18 ± 0.17 to 3.57 ± 0.08 kcal/mol. Based on the observed dwell times of nicked configurations, we designed probes for DNA-PAINT imaging and experimentally showed the applicability for simultaneous super-resolution imaging. Our work will have implications in DNA nanotechnology^6^, designing the functional nucleic acid structures^7, 8^, and, most importantly, the refinement of DNA stability predictions. In addition, the data presented in this report might aid in modeling the structure-dependent binding of proteins that require unique sequence-defined DNA topology.

### Experimental design to investigate the base-stacking interactions

We present a single-molecule imaging assay based on DNA-PAINT^25^ to deduce the base-stacking interactions between any dinucleotides. We measured the binding kinetics of fluorophore-labeled short oligonucleotide (imager strand) on its complementary strand (docking strand) using speed-optimized DNA-PAINT probes.^27, 28^ We designed two DNA configurations with the docking strand extended from a double-stranded DNA duplex (Figure 1a). In the first setup, the imager binding leaves a two-nucleotide gap between the imager’s 5’- terminal nucleotide and the 3’- recessed nucleotide of the stem (Figure 1a-top). The second configuration carries the same sequence as the first, except that it contains two additional nucleotides in place of the gap (i.e., the nick), facilitating the base-stacking interaction between the terminal nucleotides of the imager and stem (Figure 1a-bottom). These designs were inspired by the biochemical assays that attempted to quantify base-stacking interactions by gel electrophoresis of nicked and gapped DNA complexes.^3, 20^ We study the single-molecule binding kinetics of the imager to the docking strand for extracting the free energy contribution of dinucleotide base-stacking (Figure 1b).

**Figure 1:**
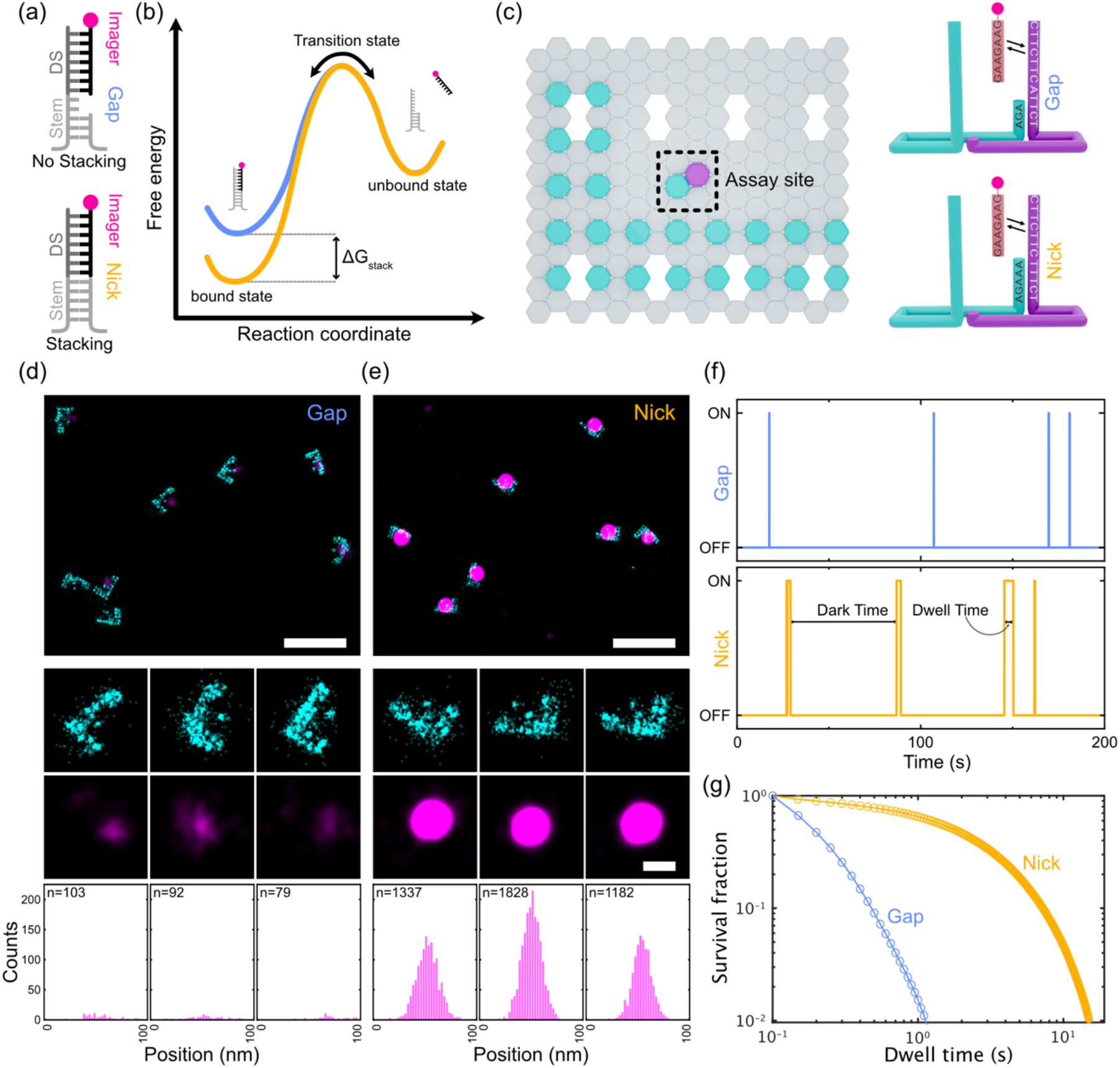
Single-molecule assay for studying dinucleotide base-stacking interactions. (a) Schematic representation of the two configurations of gap and nick. DS: Docking strand. (b) Representative free energy diagram of the bound and unbound states. Bound state under the nick configuration would show greater stabilization and thus lower free energy (orange line) than the bound state under the gap configuration (blue line). (c) Graphical representation of the origami layout. Left: The ‘L’-shaped grid with cyan-color extensions for identifying the locations of origami structures, and the assay site with magenta-color extension for studying base-stacking interactions. Right: Detailed view of the assay site where two staples are used together to generate a gap or a nick configuration. (d and e) DNA-PAINT data of origami grid imaged with ATTO647N-imager (cyan) and assay site imaged with Cy3B-imager (magenta). Gap (d) and nick (e): Row-1 showing example large field of views. Row-2 and row-3 show individual origami grids and their colocalized assay site, respectively. Row-4 shows the histograms of number of bound frames for each assay site. (f) Representative idealized individual assay site time traces indicating dark time and dwell time. (g) Cumulative survival fraction of events showing equal to or greater than shown dwell time (n= 140649 for gap and n=216040 for stack). Scale bars: 200 nm(row-1 of d, e), 40 nm (row-2 and 3 of d, e).

### Imager strand binding dynamics on the gap and nick configurations

We aimed to measure the binding dynamics of an imager strand on a docking strand with either a two-nucleotide gap or a nick at the recessed site using designer DNA origami nanostructures. For this, we designed two rectangular DNA origami structures^23^ carrying the docking strands extended from double-stranded stems in two different configurations which we call the assay site (Figure 1c). The origami structures also carry another set of docking strands in the ‘L’-shape for identifying their locations in the imaging field (henceforth called grid), enabling us to neglect spurious signals arising from any non-specific imager binding. We immobilized the origami structures on a PEG-passivated glass surface via biotin-streptavidin interactions and imaged the origami structures using two-color DNA-PAINT under a Total Internal Reflection Fluorescence (TIRF) microscope.^26, 29^ We first identified the origami structures by DNA-PAINT imaging of the origami grid using an ATTO647N-labeled imager. We then imaged the assay site using a Cy3B-labeled imager to obtain the binding kinetics under the equilibrium conditions at the gap or nick configuration (Figure 1c, Supplementary figure 1 a-d).

The reconstructed DNA-PAINT data of the gap and nick configurations showed distinct number of localizations (Figure 1d and 1e, Supplementary figure 2a). The nick configuration assay site appeared brighter, resulting from higher number of bound frames due to additional base-stacking interactions (Figure 1d and 1e-bottom). To understand the origins of this difference in bound frame numbers, we inspected the binding time traces of individual spots. The time traces showed longer dwell times on the nick configuration as compared to the gap (Figure 1f and Supplementary figure 2b). Also, the cumulative survival plot of the imager dwell times showed about ten-fold slower decay on the nick configuration (Figure 1g). These results indicated that a single extra dinucleotide base-stacking interaction substantially stabilizes the bound state of the imager, establishing that base-stacking interactions are directly measurable at the single-molecule level. Measuring absolute yet weak base-stacking energetics via this experimental approach requires comparing the binding kinetics from two different measurements, making our results sensitive to variations in ionic strength of the buffer, ambient temperature, and concentration of imager strands.^30^ This is especially relevant for our imager-docking strand hybrid as their melting point is below room temperature (21 °C).

**Figure 2:**
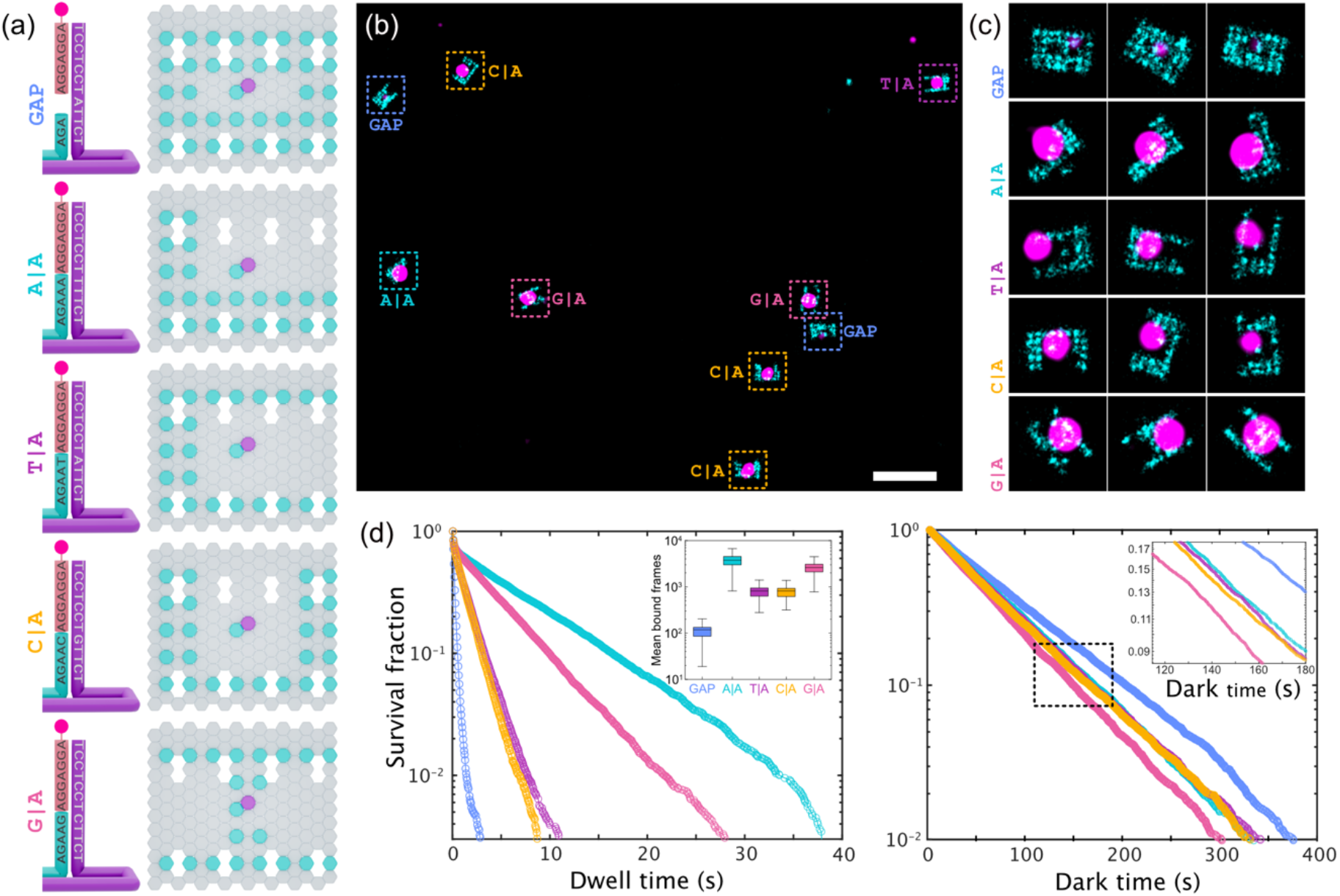
Simultaneous DNA-PAINT imaging of a gap and four base-stacking interactions. (a) Graphical representation of the five origami grids (right) and the corresponding assay sites (left) used for parallel imaging of four stack combinations possible by a single imager sequence. The stem layouts and sequences are outlined for all the five combinations where the imager stacks on top of the stem (Left). Unique grid designs enable parallel imaging (Right). (b) Representative field of view showing all five grid shapes imaged with ATTO647N-imager and corresponding assay site imaged with Cy3B-imager. (c) Representative picked origami grids and assay site. (d) Cumulative survival fractions plot showing the fraction of events at indicated dwell times (Left) or dark times (Right) greater than or equal to the given value (n= 8104, 5823, 6860, 8269, and 9425 for gap, A|A, T|A, C|A, and G|A respectively). Insert in the left side shows mean number of bound frames per assay site and right is the zoomed in view of the rectangular region. Scale bars: 200 nm (b).

### Simultaneous measurement of four base-stacking interactions

We next set up a multiplexed imaging modality to abolish any possible variations in the binding kinetics due to experimental variations. We designed five rectangular DNA origami structures carrying extensions in unique grid patterns (box-, L-, U-, C-, and H-shapes), thus making them visually distinguishable upon DNA-PAINT imaging (Figure 2a). Additionally, each grid houses an assay site consisting of a unique terminal nucleotide on the stem enabling us to image all five possible interactions (a gap and four nick configurations) with a single imager under the same conditions. For example, an imager ending with adenine nucleotide would allow us to experiment with the gap, A|A, T|A, C|A, and G|A stacking interactions (Figure 2a). Here, G|A means 5’-G stacking on 3’-A in a single-stranded DNA of 5’-GA-3’, for example.

We immobilized all five origami structures at equimolar concentrations in a microfluidic flow cell and imaged with two-color DNA-PAINT in successive rounds for identifying the grid patterns and analyzing the assay site, respectively. Reconstructed data shows clear, distinguishable origami grids, enabling us to manually identify each gap and nick configurations (Supplementary figure 3). As expected, these patterns colocalized with a single spot arising from the assay site interactions (Figure 2b). We then extracted the binding kinetics of all five configurations.

Our kinetic analysis revealed that each configuration has a characteristic dwell time distribution, indicating that the base-stacking interactions are unique to the dinucleotide combinations at the assay site (Figure 2d-left). In line with this, we also observed distinct number of bound frames for each combination (insert Figure 2d-left). Interestingly, we also observed a considerable increase in the binding frequency at nick configurations compared to the gap, evident from the dark time distributions (Figure 2d-right). These data indicate that the binding strength of the imager is dependent on the additional stabilization due to the stacked dinucleotides. We note that these data are devoid of experimental variations as they were acquired in a single imaging experiment, prompting us to directly compare the gap and nick configurations for extracting the base-stacking energetics of dinucleotides.

### Measuring all 16 possible base-stacking energetics

We next wanted to acquire binding kinetics for all 16 possible dinucleotide base-stacking configurations and deduce the corresponding free energies. We designed four separate simultaneous DNA-PAINT imaging experiments, each for extracting imager binding kinetics on four nick and a gap configuration (Inserts in figure 3a). These four imaging experiments were carried out using different imager and docking strand sequences. The DNA-PAINT experiments provided us with the imager binding kinetics. While dwell times distribution of the gap data showed a single population, the base-stacking data showed two clear populations in which shorter dwell times resembled the gap configuration, and the longer ones varied depending on the dinucleotide base-stacking under investigation (Supplementary figure 4a). This data indicates that the imager binds on nick configuration in two different modes, one with terminal nucleotides stacking and the second without stacking, equivalent to the gap (Supplementary figure 4b). The appearance of unstacked population is likely due to the fraying of the stem. This hypothesis was corroborated by the fact that we observed several blocks of consecutive short-lived and blocks of long-lived binding events. If the stem was always in the hybridized state, we would not have expected this segregated consecutive short-lived and long-lived binding events (Supplementary figure 4b). Based on this, we built a kinetic model for both the gap and nick configurations for extracting the rate constants for imager binding (Supplementary figure 4c, see Supplementary methods).

**Figure 3:**
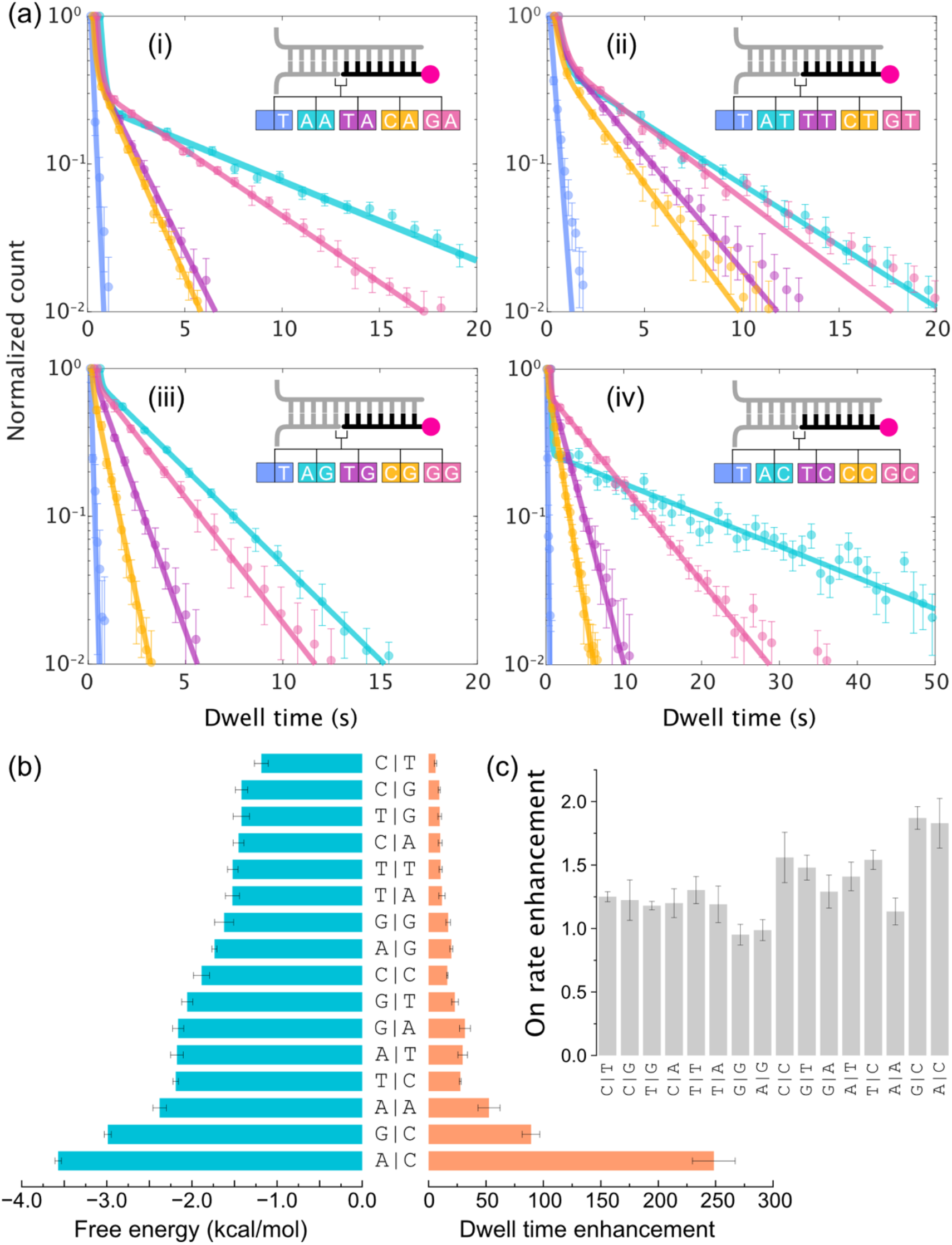
Imaging base-stacking interactions of all 16 possible dinucleotide combinations. (a) Histograms showing dwell time distributions (points) and mathematical fits to the data (curves). Each plot obtained from single simultaneous imaging rounds, show four base-stacking interactions and a gap as represented in the insets with the corresponding color combinations. All the gap data sets were fit with mono-exponential and nick data sets were fit with a bi-exponential function to obtain corresponding off-rate constants. Error bars were generated via bootstrapping analysis. (b) The dwell time enhancement obtained by taking the ratio of off-rate constants of gap to that of the nick (right), and the calculated dinucleotide base-stacking free energetics (left). (c) On rate enhancement obtained from calculating the ratio of binding rate on gap to nick.

The gap dwell time histogram data was fit with a mono-exponential function providing us with the off-rate constant (*k*_*off*_) (Figure 3a and Supplementary figure 4d). The nick data were fit with the bi-exponential function, which provided two off-rate constants in which the first one (*k*_*off*,1_) was evidently similar to the gap configuration (i.e., *k*_*off*_), and the second one (*k*_*off*,2_) was characteristic of the nick configuration (Figure 3a and Supplementary Figure 4c, 4d). This statistical analysis of dwell time distribution provided us the actual off-rate constants (*k*_*off*,2_) for the corresponding imagers on all 16 nick and four gap configurations (Supplementary table 1). Next, we calculated the fractional enhancement in the binding time by taking the ratio of the gap to nick configuration (*i*.*e*., 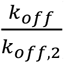) which showed a wide distribution depending on the dinucleotide combination, starting from around 6-fold for C|T to 250-fold for A|C (Figure 3b). We suspected the unexpectedly large fold change in the binding time was due to the sequence context. We challenged this surprisingly large fractional enhancement for A|C by designing an orthogonal assay site configuration with a different stem sequence. We observed similar fractional enhancement in dwell times validating the accuracy of our experimental approach (Supplementary figures 5a and 5b). This data not only confirms our observation that the A|C is the strongest interacting dinucleotide but also suggests that the sequence context of the stem may not substantially contribute to the base-stacking interactions of the dinucleotides under investigation. Additionally, we observed identical fractional enhancement when we substituted our experiment with a different fluorophore on the imager strand (Supplementary figure 5c), ruling out the possibility of the fluorophore intervention in the measured base-stacking kinetics. Fluorophore photobleaching presents a limitation for the single-molecule imaging experiments resulting in the underestimation of actual binding times.^31^ However, we note that the fluorophore photobleaching rate (*k*_*photobleaching*_= 0.0007 sec^−1^) is substantially lower than the slowest measured *k*_*off*,2_ = 0.043 sec^−1^ for A|C (Supplementary figure 6) in our experimental conditions, indicating that these measurements are not suffering from the photo-physical properties of the fluorophore.

The DNA-PAINT data also provides us with the dark times between the consecutive imager binding events on the docking strand. The dark times histogram showed a clear mono-exponential distribution. We extracted the binding frequency (*k*_*bind*_) by fitting the corresponding histogram with a mono-exponential function. In most cases, the nick configurations showed a considerable increase in the *k*_*bind*_ compared to gaps (Figure 3c, Supplementary figure 7, and Supplementary table 2). We anticipate the nick configuration would mechanistically have two opposing effects on *k*_*bind*_. First, the imager would experience a steric hindrance by the stem site, which would negatively impact the initiation of binding, hence would result in decreased *k*_*bind*_. Second, base-stacking interaction between the recessed nucleotide on the stem and the terminal nucleotides of the imager acts as an additional nucleation site for binding, resulting in increased *k*_*bind*_.^32^ Indeed, we observe a positive correlation between the dinucleotide stacks with fractional enhancement in dwell time and enhancement in *k*_*bind*_ (Supplementary figure 8a). Additionally, enhancement in the *k*_*bind*_ negatively correlated with the bulkiness of the underlined stacked dinucleotides that could potentially cause steric hindrance for the imager binding (Supplementary figure 8b). The overall increase in the *k*_*bind*_ is likely because of base-stacking interactions outcompeting the steric hindrance.^32^

We ultimately wanted to calculate the absolute base-stacking free energies for all 16 combinations based on our single-molecule kinetic data. By comparing the binding kinetics of imager on stacking configuration and gap (see Supplementary Methods), we provide the absolute base-stacking free energy for each dinucleotide combination (Figure 3b).

The overall trend is that the base-stacking energetics presented here correspond to the degree of molecular overlap of nitrogenous aromatic rings in the dinucleotide combination (Supplementary figure 9). This fact is substantiated when comparing the swapped-sequence pairs that have the same molecular composition, such as A|C and C|A, G|C and C|G, etc., but show distinct interactions because they interact via dissimilar exposed molecular surfaces (Supplementary figure 9). The reverse complement dinucleotides, such as A|C and G|T, C|T and A|G, etc., also showed rather distinct interactions energies owing to the characteristic molecular interactions. This surprising observation was only possible as we could disentangle the individual dinucleotide base-stacking interactions rather than measuring complete base-pair stacking interactions. ^18, 22^ Intriguingly, the average of our individually measured reverse complement sequence energetics are in close agreement with the previously reported base-pair stacking energetics (see Supplementary table e).^18, 22, 33, 34^

Our data is in qualitative agreement with the previous results showing purine|pyrimidine interactions are in general stable than their counterparts^11^, but the absolute free energies are surprisingly higher.^3, 18, 22, 33^ This discrepancy is likely due to varied experimental strategies and conditions used across different studies. More importantly, the current report presents the direct single-molecule measurement of the individual base-stacking interactions, unlike the previous reports that estimated base-pair stacking interactions.^18, 22, 33, 35^

### Stack-PAINT for simultaneous multiplexed super-resolution imaging

We next wanted to test the applicability of the imager’s binding kinetics dependency on stacking interactions for simultaneous multiplexing in DNA-PAINT super-resolution imaging.^36^ We envisioned “three-color” simultaneous multiplexing based on the “tunable” imager binding dwell times to its cognate docking strand depending on the underlined stacking interactions (Supplementary Figure 10a-10d). For a proof-of-principle demonstration, we chose an imager strand with adenine terminal-nucleotide and three different docking strands in which one does not form a stem and two others form a stem with either adenine or thymine at the termini (i.e., gap, A|A, and T|A). On these three configurations, the imager strands are expected to bind on average around 150 ms, 1.5 s, and 8 s, respectively (Supplementary Figure 10a). We designed three DNA origami grids with docking strand extensions including these stem configurations (Figure 4a). The docking strands were separated by varying distances starting from 20 nm for testing the super-resolvability of multiplexed DNA-PAINT. As the binding kinetics are engineered based on the stacking nucleotides, we termed this modality Stack-PAINT. For ground truth verification, we also decorated the origami structures with another docking strand in specific grid patterns (‘H’-, ‘C’-, and ‘U’-grids) (Figure 4a).

**Figure 4:**
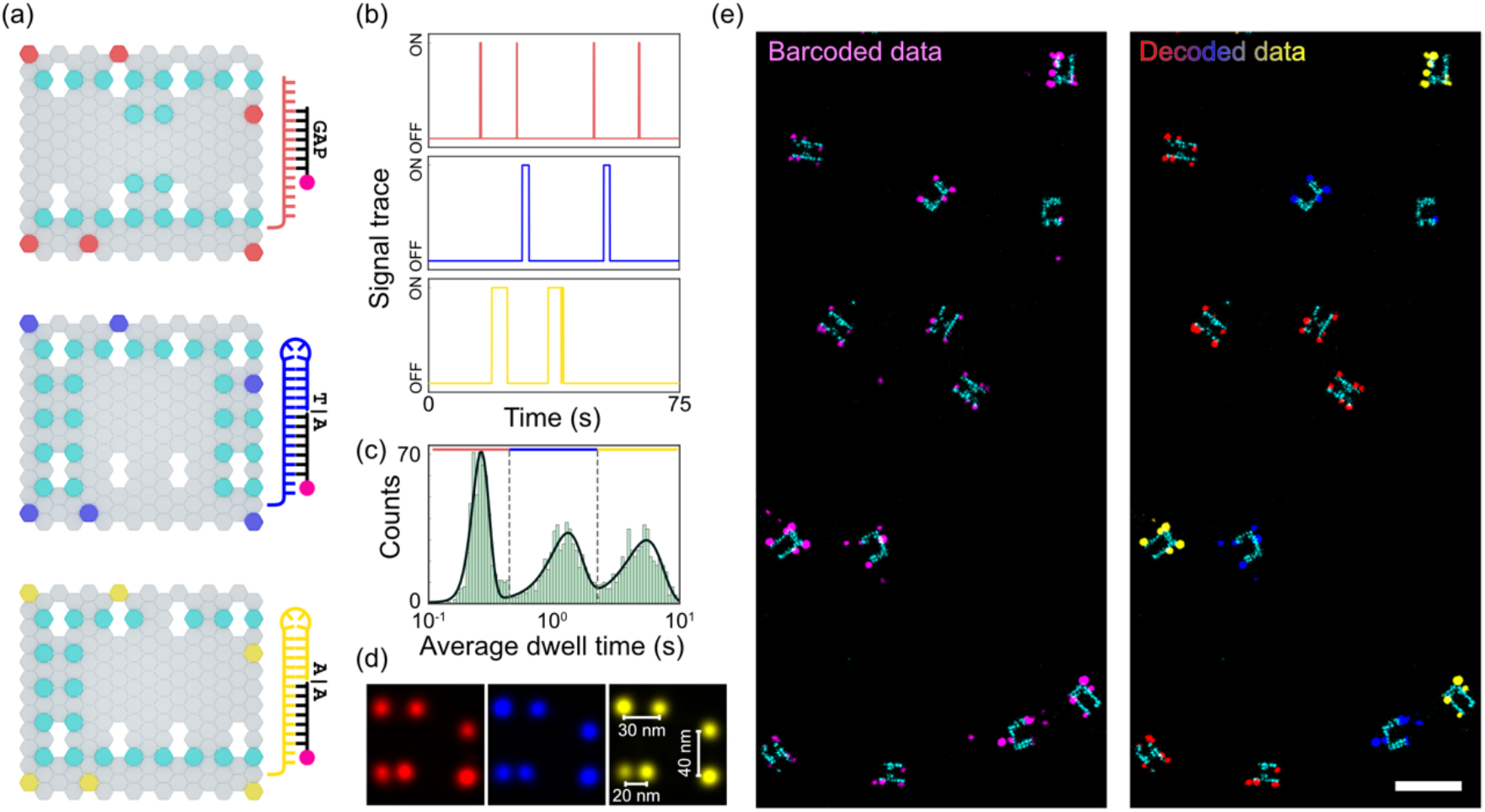
Stack-PAINT: Application of base-stacking energetics for multiplexed simultaneous super-resolution imaging. a) Schematics of origami structures for multiplexed DNA-PAINT imaging. Cyan extensions are for ground truth identification. The design details of the colored extensions are shown on the right side. Gap (Red), T|A (Blue) and A|A (Yellow)are bound by the same imager with different binding strength because of the different stacking or gap interactions, facilitating multiplexed imaging. b) Representative idealized imager binding time traces. c) Histogram showing average dwell time distributions from all the origami structures in the field of view (n=1463). The dotted lines demarcate the peaks based on the expected average binding times and the colored lines represent the gap or stack data. Color code is similar to (a) d) Overlaid DNA-PAINT origami structures taken from the demarcated histogram data. The color code is similar as stated in (a). (n=525, n=456, and n= 425 in the order of images). e) Representative barcoded DNA-PAINT data (left) and decoded data (right) based on the average dwell times corresponding to the ground truth (cyan). The color code is similar as stated in (a).

Upon Stack-PAINT imaging, individual time traces of imager bindings showed the expected dwell times for each of the nick configuration (Figure 4b). We obtained the average dwell times of the imager on individual origami structures in the entire field of view. The histogram of the average dwell times resulted in three distinguishable populations as expected, each originating from three different configurations (Figure 4c). We accordingly classified the origami structures under each peak into separate populations (Figure 4c). The origami structures were then transformed into pseudo-colored, barcoded images based on the matching dwell times. We ascertained that Stack-PAINT could resolve 20 nm separated docking strands using all three configurations (Figure 4d). We matched the kinetically analyzed Stack-PAINT data with the ground truth grid patterns with high accuracy (∼97%) (Figure 4e and Supplementary Figure 11), demonstrating the applicability of specific stacking interactions for multiplexed super-resolution imaging. In similar lines, we envision that our stacking energetics data will enable the design of novel DNA-PAINT probes with tunable kinetics.

## CONCLUSIONS

This work presents direct measurement of a comprehensive list of dinucleotide base-stacking interactions providing all 16 combinations. We developed a multiplexed, high-throughput single-molecule imaging assay exploiting the power of DNA nanotechnology to extract the binding kinetics of fluorophore-labeled short single-stranded DNA. As DNA-PAINT records the transient interactions of repetitive imager binding on the docking strand under equilibrium, we obtained about half a million molecular hybridization events which is required for robust statistical analysis of binding kinetics,^37^ thereby facilitating the deduction of the absolute dinucleotide base-stacking energetics with high accuracy of ±0.1 kcal/mol. We carefully analyzed the kinetics under orthogonal configurations to ascertain our measured values with high confidence.

Given that the base-stacking interactions provide great control over the imager binding times, we exploited this engineered kinetics for the designing of DNA-PAINT imaging probes for multiplexed high-resolution microscopy. The novel design enabled for multiplexed imaging while preserving the super-resolution aspect of DNA-PAINT. In combination to the sequence composition of the imager and salinity of the buffer, base-stacking energetics provide multifaceted control over the binding kinetics of the imager strands and will drive the development of novel DNA-PAINT probes. This result might inspire for designing of RNA and DNA aptamers for small molecule sensing. As the multiplexed imaging strategy described in this study only requires a widely used TIRF microscope without any external mechanical manipulation of molecules, it can readily be extended to study other nucleic acid interactions including RNA, PNA, and LNA that might contain different chemically modified nucleotides.

## Author Contributions

A.B. conceived and performed experiments, analyzed data, and wrote the manuscript. M.A. analyzed and curated data. S.K. performed experiments. M.G. conceived and supervised the study, analyzed and interpreted data, and wrote the manuscript. All authors reviewed and approved the manuscript.

## ACKNOWLEDGMENT

We thank Dr. Thomas Schlichthaerle, Dr. Sung Hyun Kim, Dr. Rafayel Petrosyan, Prof. Harinder Singh, and Dr. Sandeep Choubey for fruitful discussions. We thank Dr. Saminathan Ramakrishnan for the preliminary experiments. We greatly acknowledge Dr. Sarit Agasti for allowing us to use their laboratory facilities. A.B. acknowledges the support from the Prime Minister’s Research Fellowship (PMRF), Ministry of Education, Government of India. M.A. and S.K. acknowledge the support received from Council for Scientific and Industrial Research, Ministry of Science and Technology, Government of India. We acknowledge the Department of Science and Technology, Ministry of Science and Technology, India DST-FIST Program funded Central Facility, Department of Biochemistry, IISc.

