## Supplementary material for "Single-Molecule Analysis of DNA Base-Stacking Energetics Using Patterned DNA Nanostructures"

#### Methods

##### Slide preparation

Microscopy slide and coverslip preparation was done as previously described.[1,2] The slides (VWR #631-1550) were drilled using a diamond head drill bit (Meisinger #801-009-HP) and drill gun (DigitalCraft #LRUXOR). Five holes on each side spaced about 5 mm apart were made to allow for preparation of 5 microfluidic channels on each slide. In case the slides were being reused, macroscopic particles were removed using 5% v/v dish washing detergent before further steps were performed. Slides and coverslips VWR #631-0147) were initially rinsed thoroughly using MilliQ water and then immersed into coplin jars (Tarsons #480000) containing MilliQ water. The slides and coverslips were then sonicated for 5 minutes (IGene LabServe #IGNUC-9). MilliQ water was then replaced with Acetone (SRL #31566) twice and the slides and coverslips were sonicated for 5 minutes. Acetone was then replaced with MilliQ water followed by 1 M KOH (BDH #296228) and sonicated for at least 30 minutes. The slides and coverslips were then rinsed using MilliQ water by replacing the KOH in the coplin jars at least thrice. The slides and coverslips were then sonicated in MilliQ water for 10 minutes and dried using compressed nitrogen gas.

Piranha etching was performed on the slides and coverslips. The slides and coverslips were placed in dried coplin jars. Piranha solution was prepared by adding one part of 30% $H_2O_2$ (EMPLURA #107209) to three parts of $H_2SO_4$ (Fisher Scientific #29997) and stirred well until boiling. This solution was then transferred into the coplin jars and left for 30 minutes. The piranha solution was disposed of in a dedicated waste container and the slides and coverslips were rinsed thoroughly using MilliQ water. The slides and cover slips were then rinsed with methanol twice and were sonicated in methanol for 20 minutes.

Aminosilanation was then performed after thorough cleaning and etching of the slides. A dedicated container for aminosilane mixture preparation was made by sonicating the container filled with methanol. The container can be reused repeatedly if it is refrigerated with methanol in a sealed condition using parafilm. The aminosilanation mix was prepared by mixing 5 ml of acetic acid (SDFCL #20001) in 100 ml of methanol, followed by addition of 10 ml of APTES ((3-Aminopropyl) triethoxysilane) (SRL #33993 or TCI #A0439) and mixed thoroughly. This solution was then poured over the slides and coverslips held within the coplin jar and left undisturbed for 25 minutes. The slides and coverslips were washed thoroughly with fresh methanol three times followed by rinsing with MilliQ water. Slides were then dried with compressed nitrogen gas.

Glass slides and cover slips were passivated by covalent modification with PEG. Passivation was performed on these aminosilanated slides using mPEG-SVA (Succinimidyl Valerate) (Laysan Bio #mPEG-SVA-5000) and Biotin-PEG-SVA (Laysan Bio #Biotin-PEG-SVA-5000). mPEG-SVA and Biotin-PEG-SVA were dissolved in a ratio of 40:1 by mass in 0.1M $NaHCO_3$ (Sigma #S5761) pH 8.5. For 15 pairs of slides and coverslips, 120 mg of mPEG-SVA and 3 mg of Biotin-PEG-SVA was dissolved in 960 $\mu$l of 0.1 M $NaHCO_3$. 60 $\mu$l of this solution was added onto each slide and then sandwiched by placing a coverslip over it gently. Trapped air bubbles escape over the incubation period. These slides and coverslips were stored in humid chambers made by pouring some water into empty micropipette tip boxes. These boxes were then stored in dark for a duration of eight to twelve hours at room temperature. The following day, the sandwich was disassembled and washed thoroughly with MilliQ water. These slides were then dried with compressed nitrogen gas and stored inside 50 ml centrifuge tubes (NUNC #339653) with the PEG-passivated surfaces facing away from each other. These tubes were vacated using a vacuum pump and stored under inert conditions in nitrogen gas. The lids of these tubes were wrapped with parafilm and stored in a freezer for later use.

The flow cells were assembled using double sided tape (3M Scotch #136D MDEU) between each pair of holes on the PEGylated sides. Coverslips were placed carefully of the taped slide ensuring that the PEG-passivated slides make up the interiors of the microfluidic flow cells. The open edges of these channels were sealed using epoxy.

DNA Origami Folding

The rectangular DNA origami structure was taken as the template for the origami structures used in this study. We used the Picasso design module[3] to get the staple sequences of specific extensions for the grids and blank staples (Supplementary table 4 and 5). M13mp18 ssDNA (Bayou Biolabs #P107) was used as the scaffold for the DNA origami. Biotinylated oligonucleotide staples (Supplementary table 6) were used to anchor the origami structure to the flow cell surface. Staples with extension for the gap and stack positions were designed using sequences from Picasso design and CadNanoSQ (https://cadnano.org/, Supplementary table 7). Folding Buffer contained 50 mM Tris-Cl (Tris-Base, Sigma #77861; HCl, Fisher Scientific #29507) pH 8.0, 12.5 mM $MgCl_2$ (EMPLURA #105833) and 0.2 mM EDTA (SRL #35888). We set up 30 $\mu$l reactions with 10 nM of scaffold DNA, 100 nM of biotin staples, 100 nM of blank staples, 1 $\mu$M of staples for the grid and 33.3 $\mu$M of gap or stack specific staple in the folding buffer. The mix was then heated up to 80 °C, held for five minutes, and then slowly cooled to 4 °C, in steps of 0.1 °C every five seconds in a thermocycler.

The folded origami structures were purified using Sartorius Vivaspin® 500 (Sartorius #VS0132) centrifugal filters. Equilibration was performed by spinning the columns at 3000 ×g for 5 minutes with 500 $\mu$l of HPLC grade water (SRL #92605) followed by 500 $\mu$l of folding buffer. The resultant origami mix was then applied to the column along with 500 $\mu$l of folding buffer at 800-1000 ×g for three rounds, to remove most of the unincorporated staples. Origami structures were stored in the folding buffer at a concentration of 3.3 nM at -20 °C.

Imager fluorophore conjugation and purification

We obtained 3'-end amine modified oligonucleotides from Sigma (Supplementary table 8) and dissolved to a final concentration of 1 mM using HPLC grade water.

Fluorophores were dissolved in DMSO (Sigma #D8418) to the following concentrations. Cy3B-MonoNHS-Ester (Cytiva #PA63101) was dissolved to a final concentration of 13 mM, Atto647N-MonoNHS-Ester (Sigma #18373-1MG-F) was dissolved to a final concentration of 11.8 mM and Cy5-MonoNHS-Ester (Cytiva #PA15101) was dissolved to a final concentration of 1.3 mM.

A 10× PBS was prepared by adding 80 g of NaCl (SRL #33205), 2 g of KCl (Sigma #P9541-1KG), 14.4 g of $Na_2HPO_4$ (SRL #1949146) and 2.4 g of $KH_2PO_4$ (Ranbaxy #5HEV0740) in one liter of MilliQ water followed by autoclaving at 121 °C for 15 minutes. The pH of the buffer was adjusted to 7.4.

For conjugation of Cy3B or Atto647N to the imager, 15 nanomole of DNA (15 $\mu$l from 1 mM stock) was dissolved in a mixture containing 3 $\mu$l 10× PBS, 3 $\mu$l of 1M $NaHCO_3$ and 3.24 $\mu$l HPLC grade water. 75 nanomole (5.76 $\mu$l of 13 mM Cy3B or 6.35 $\mu$l of 11.8 mM Atto647N) of fluorophore was added to the above mixture and vortexed vigorously. The mix was incubated overnight in the dark at room temperature under vigorous shaking.

For conjugation of Cy5 to the imager, 2 nanomole of DNA (2 $\mu$l from 1 mM stock) was dissolved in a mixture containing 1.5 $\mu$l 10× PBS, 1.5 $\mu$l of 1 M $NaHCO_3$ and 4 $\mu$l HPLC grade water. 7.8 nanomole (6 $\mu$l of 1.3 mM Cy5) of fluorophore was added to the above mixture and vortexed vigorously. The mix was incubated overnight in the dark at room temperature under vigorous shaking.

Post overnight incubation, the conjugated product was purified from the free fluorophore and unconjugated DNA oligonucleotide using reverse phase (Phenomenex # 00B-4442-E0 Clarity® 5 $\mu$m Oligo-RP, LC Column 50 × 4.6 mm, Ea) HPLC (Agilent technologies). The conjugated product was then dissolved in HPLC grade water and stored at -20 °C until using.

Sample preparation

Microfluidic flow cells with PEG-passivated cover slip and slide were incubated with 10 $\mu$l of 0.2 mg/ml Neutravidin (Sigma #31000) in T50 buffer containing 50 mM Tris-Cl pH 8.0, 50 mM NaCl and 0.2 mM EDTA for 20 minutes. This was followed by thorough washing of the microfluidic channel with 600 $\mu$l of T50 buffer. Imaging/Immobilization buffer (Buffer I) containing 50 mM Tri-Cl pH 8.0, 10 mM $MgCl_2$ and 0.2 mM EDTA was used to wash the channel prior to origami immobilization. The origamis intended for a specific imaging run were pooled together at a final concentration of 400-600 pM each in buffer I and applied onto the channel and incubated for 20 minutes. This was followed by washing of the channel with 600 $\mu$l of buffer I prior to imaging to remove any unbound origamis.

Microscopy and Imaging

Microscopy was performed on a Nikon Ti2 eclipse microscope equipped with a motorized H-TIRF, perfect focus system (PFS), and a Teledyne Photometrics PRIME BSI sCMOS camera. Illumination using 561 nm and 640 nm wavelength lasers was done using the L6cc Laser combiner from Oxxius Inc., France. Imaging was done under Total Internal Reflection conditions. An oil immersion objective lens (Nikon Instruments #Apo SR TIRF 100×, NA 1.49, oil) was used for imaging. Imaging was performed with 2 × 2 binning of pixels and the camera was cropped to an effective size of 512 × 512 pixels, each pixel spanning 130 × 130 nm. Acquisition was done by setting the camera to a readout sensitivity of 16-bit. Imaging parameters used in the different experiments are outlined in Supplementary table 14.

20× PCD was made by dissolving PCD (Protocatechuate 3,4-dioxygenase; Sigma #P8279-25UN) in buffer containing stock in 50 mM KCl, 1 mM EDTA and 100 mM Tris–HCl, pH 8.0 and 50% glycerol (Sigma #G5516-1L) to a final concentration of 6 $\mu$M. The solution was divided into 10 $\mu$l aliquots in PCR tubes and stored at -20 °C.

40× PCA was made by dissolving 154 mg of PCA (Protocatechuic acid/ 3,4-Dihydroxybenzoic acid; Sigma #37580-100G-F) in 10 ml of HPLC grade water adjusted to pH 9.0 using 1 M NaOH (SRL #96311). The solution was divided into 100 $\mu$l aliquots in 200 $\mu$l tubes and stored at -20 °C.

100× Trolox was prepared by dissolving 100 mg of Trolox (6-hydroxy-2,5,7,8-tetramethylchroman-2-carboxylic acid; Sigma #238813-1G) in 430 $\mu$l of methanol, 345 $\mu$l of 1 M NaOH and 3.2 ml of HPLC grade water. The solution was divided into 20 $\mu$l aliquots in PCR tubes and stored at -20 °C.

Prior to imaging, 100 $\mu$l of imaging buffer was prepared in buffer I with a final concentration of 1× PCA, 1× PCD and 1× Trolox, along with imagers at concentrations mentioned in Supplementary table

14. Imaging was always performed with the 640 nm laser prior to the 561 nm laser to minimize the effect of ATTO647N or Cy5 fluorophore photobleaching by the higher energy (561 nm) light source.

Imaging of stem-loop layout (Supplementary figure 5)

Origamis were folded in the same manner as mentioned above (Supplementary table 4, 5, 9, and 11). Imaging was performed using Exchange-PAINT[4], where first round of imaging was done to image the origamis carrying R1×5 extensions followed by origamis carrying R4×5 extensions with appropriate imagers carrying Atto647N fluorophore at mentioned concentrations (Supplementary table 14). Imaging buffer was prepared as mentioned above. Subsequent imaging rounds were separated by washing with 2 ml of imaging buffer. Finally, the stack and gap were measured using Cy3B-labelled imager at the concentrations mentioned in the Supplementary table 14.

Data Analysis

The obtained raw fluorescence data was reconstructed using 'Picasso Localize' software package[3] to obtain a super-resolved image. Drift correction was performed by redundant cross-correlation (RCC). RCC was also used to align the super-resolved structures from the two imaging channels (Supplementary Figure 1). Origamis were manually picked based on their grid structures. Localizations from the gap or nick spots from the assay sites were extracted using 'Picasso Render' for further kinetics analysis. We performed kinetic analysis using a custom written MATLAB code. Briefly, the list of localizations exported for each origami pick was further analyzed for individual dwell times and dark times. Dwell times were calculated based on the presence of consecutive binding events with a gap of not more than 25 frames (1250 milliseconds at an acquisition rate of 20 frames per second with an exposure time of 50 milliseconds) (see figure 1f). This is done to overcome any potential flickering of the fluorophore and the lower signal-to-noise-ratios that arise due to the lower laser powers that were used during imaging to maintain the photostability of the fluorophore (Supplementary Figure 6). Further, we are confident that this will not combine two consecutive binding events due to the extremely long-time intervals of around 100 to 200 seconds between consecutive bindings (Supplementary Figure 7). Bootstrapping for 100 iterations was performed based on the mean using MATLAB's built-in bootstrap function on the list of individual dwell times for each data set separately.

Dark times were calculated based on the durations between two consecutive binding events (see figure 1f). The obtained dark times were bootstrapped with parameters mentioned above and used for further kinetic analysis.

Kinetic analysis for calculating the Base-stacking energies

The gap configuration is modeled with a bound state and an unbound state with corresponding binding rate ($k_{bind,gap}$) and dissociation rate ($k_{off}$). The bound state in the nick configuration consists of two subpopulations (i.e., the bound-stacked state and bound-unstacked state) characterized by two dissociation rate constants, the rate of dissociation from the bound-unstacked state and the bound-stacked state to unbound state ($k_{off,1}, k_{off,2}$) and a binding rates ($k_{bind,nick}$). kinetics analysis facilitated us calculating the dissociation constant ($K_D$) which provides us with the free energy ($\Delta G$) of the duplex formation on each substrate. We used following formalism to extract the base-stacking free energy contribution of individual dinucleotides.

$$Free\ energy\ for\ gap/nick\ configuration, \Delta G = -R \cdot T \cdot ln(K_D)$$

$$where, \qquad K_D = \left(\frac{C \cdot k_{off}}{k_{bind}}\right)$$

$$\Delta G_{nick} = -R \cdot T \cdot ln\left(\frac{C \cdot k_{off,2}}{k_{bind_{nick}}}\right)$$

$$\Delta G_{gap} = -R \cdot T \cdot ln\left(\frac{C \cdot k_{off}}{k_{bind_{gap}}}\right)$$

$$Base - stacking\ free\ energy, \qquad \Delta G_{stack} = \Delta G_{nick} - \Delta G_{gap}$$

Here, $\Delta G_{nick}$ represents the free energy of imager binding on the nick configuration and, $\Delta G_{gap}$ represents the free energy of imager binding on the gap configuration. As described above, we assessed the frequency of binding ($k_{bind}$) and off-rate ($k_{off}$) from the kinetic analysis of the single-molecule data to estimate the free energy of duplex formation with both the gap and nick configuration. Here $C$ is the concentration of imager used in the experiment. Note that the $\Delta G_{stack}$ is independent of the imager concentration as the gap and nick configurations are measured in the same imaging experiments. The absolute temperature $T$ is constant. If the $T$, $C$, and the salinity of the buffer in independent experiments are not well controlled, larger variations are expected in the free energy estimations as the binding dynamics of short oligonucleotides are strongly dependent on these parameters.[5] The multiplexed experiments utilized in the current study are immune to such possible variations as the normalization is internal to individual imaging experiments.

Here we describe the kinetic analysis of the single-molecule imaging data. The set of bootstrapped dwell times were plotted on histogram and then fit with a mono-exponential curve ($y = a_1 \cdot e^{k_{off} \cdot t}$) in case of gap, resulting in the off-rate constant $k_{off,1}$. For the nick configurations data, we fit them with a biexponential curve ($y = a_1 \cdot e^{k_{off,1} \cdot t} + a_2 \cdot e^{k_{off,2} \cdot t}$). Distribution of individual dwell times displayed in Supplementary Figure 4a clearly shows that the imager binds on the docking strand in two modes – one without the terminal nucleotide stacking on the stem, leading to a faster decaying population, and the second with terminal nucleotides stacking, leading to a slower decaying population away from the first population (Supplementary figure 4a). As we observed two distinct populations in the individual dwell time distributions, we assumed that the unstacked binding mode was due to the stem undergoing fraying temporarily. The fraying of the stem is mechanistically equivalent of the gap construct. As a result, we ignored transitions between the stack and unstacked configurations in a binding event and depicted the kinetic model in supplementary figure 4b. This fitting results in two off-rate constants ($k_{off_{,1}}$ $and$ $k_{off_{,2}}$) in which $k_{off_{,1}}$denotes the dissociation from unstacked-bound state to the unbound state which closely resembles the gap off-rate constant. The second off-rate constant, $k_{off_{,2}}$, depicts the imager dissociation rate from the stacked-bound state to the unbound state. This kinetic analysis provides with the off-rate constants of unstacked- and stacked-bound state based on the experimentally measured dwell times.

In the similar manner, we also obtained the binding rate constant ($k_{bind}$) by kinetic analysis of dark times distributions. We built histogram of the bootstrapped dark times and fit with a mono-exponential curve ($y = b_1 \cdot e^{k_{bind} \cdot t}$) to obtain the individual binding rate constants for the gap and nick configurations.

We then proceeded with obtaining the free energy of dinucleotide base-stacking interactions using the kinetically derived $k_{bind_{gap}}, k_{bind_{nick}}, k_{off}$ $and$ $k_{off_{,2}}$.

<u>Photobleaching calculation</u>

Origamis carrying the S1 sequence (Supplementary table 12) were folded using methods mentioned above. DNA origamis were immobilized on flow cells treated with neutravidin as mentioned above. Complimentary S1 strand carrying Cy3B was flown in at 1 pM concentration in buffer I. After 10 minutes of incubation, the flow cells were washed with 1 ml of buffer I to remove any unbound complimentary S1 strands carrying Cy3B. Imaging buffer was prepared with 1× PCA, 1× PCD and 1× Trolox. For recreating the exact same conditions of the Cy3B imaging round (second imaging round), the imaging buffer was added to the flow channel and a dummy imaging run was performed for the duration of the first imaging rounds. This was followed by imaging of the stably bound Cy3B with 561 nm laser excitation in TIRF mode at a similar power to the experiments depicted in figure 3 and mentioned in Supplementary table 14.

The acquired image was run through the 'Picasso Localize'[3] package to track individual fluorophores over a time course. Individual photobleaching times were obtained from the time traces generated similar to image analysis described above. These individual bleaching times were then plotted in a histogram. The fluorophores surviving throughout the imaging run and a small population of fluorophores that bleach during the first 100 seconds (first bin in the supplementary figure 6) were ignored for exponential fitting. The plotted events were then fit with a mono-exponential curve ($y = a \cdot e^{k_{photobleach} \cdot t}$) to obtain the photobleaching rate.

Stack-PAINT imaging

Origami structures were folded with staples defined in supplementary table 13 and purified using the centrifugal filtration. The origami samples were immobilized on the surface of PEG-passivated glass slide as described above. Imaging was performed as mentioned in supplementary table 14. Reconstructed data was aligned for both channels. Origamis were picked based on the six extensions that were placed for resolution testing and then filtered based on the presence of the underlying grid. The mean dwell times at each picked origami structure was calculated. We then constructed a histogram from the mean dwell times and fitted with triple-gaussian curve using Matlab 'gauss3' function. Each of the three gaussian peaks were split into three data sets by manual demarcation based on the known dwell time averages over each pick. For assessing robustness of our decoding technique, we manually filtered the selected origami structures based on the grid structures. We then compared the manual selection set to the set delineated based on the average dwell times. We calculated the error in calling the correct origami by taking the ratio of origami numbers that fall outside the expected region to total origami structures analyzed.

### Supplementary figures and captions

Supplementary Figure 1

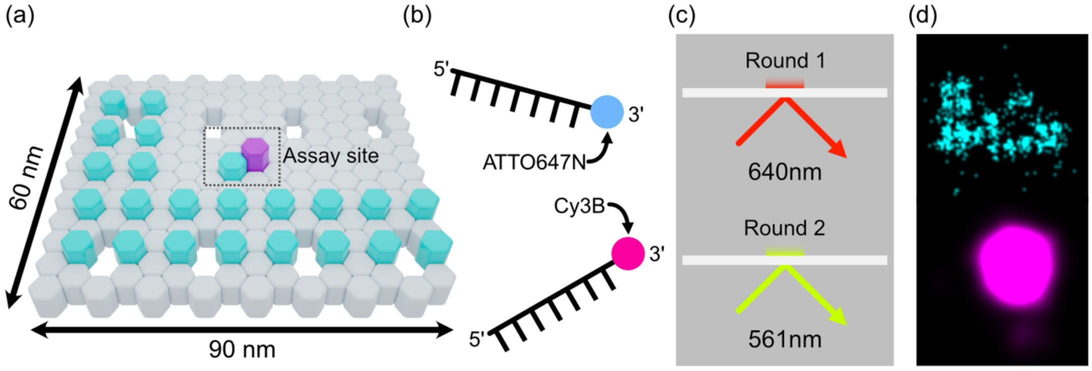


**Supplementary Figure 1:** DNA origami design and imaging scheme. (a) Graphical layout of the DNA origami design used in the experiments. Cyan hexagons represent the extensions used to image the grid. Magenta hexagon is used to represent extension at the assay site. (b) Graphical representation of the two fluorescent imager strands used in the experiments. Imager strand labelled with ATTO647N (top) was used for exciting with the 640 nm laser, and imager labelled with Cy3B (bottom) was used with the 561 nm laser. (c) Imaging was performed in TIRF mode in two rounds. First, using the 640 nm laser to excite the ATTO647N fluorophore-labelled imager for imaging the grid (cyan in (a)), and second, using the 561 nm laser to excite the Cy3B fluorophore-labelled imager for imaging assay site (magenta in (a)). (d) Example super-resolved images obtained from the two rounds of imaging.

Supplementary Figure 2

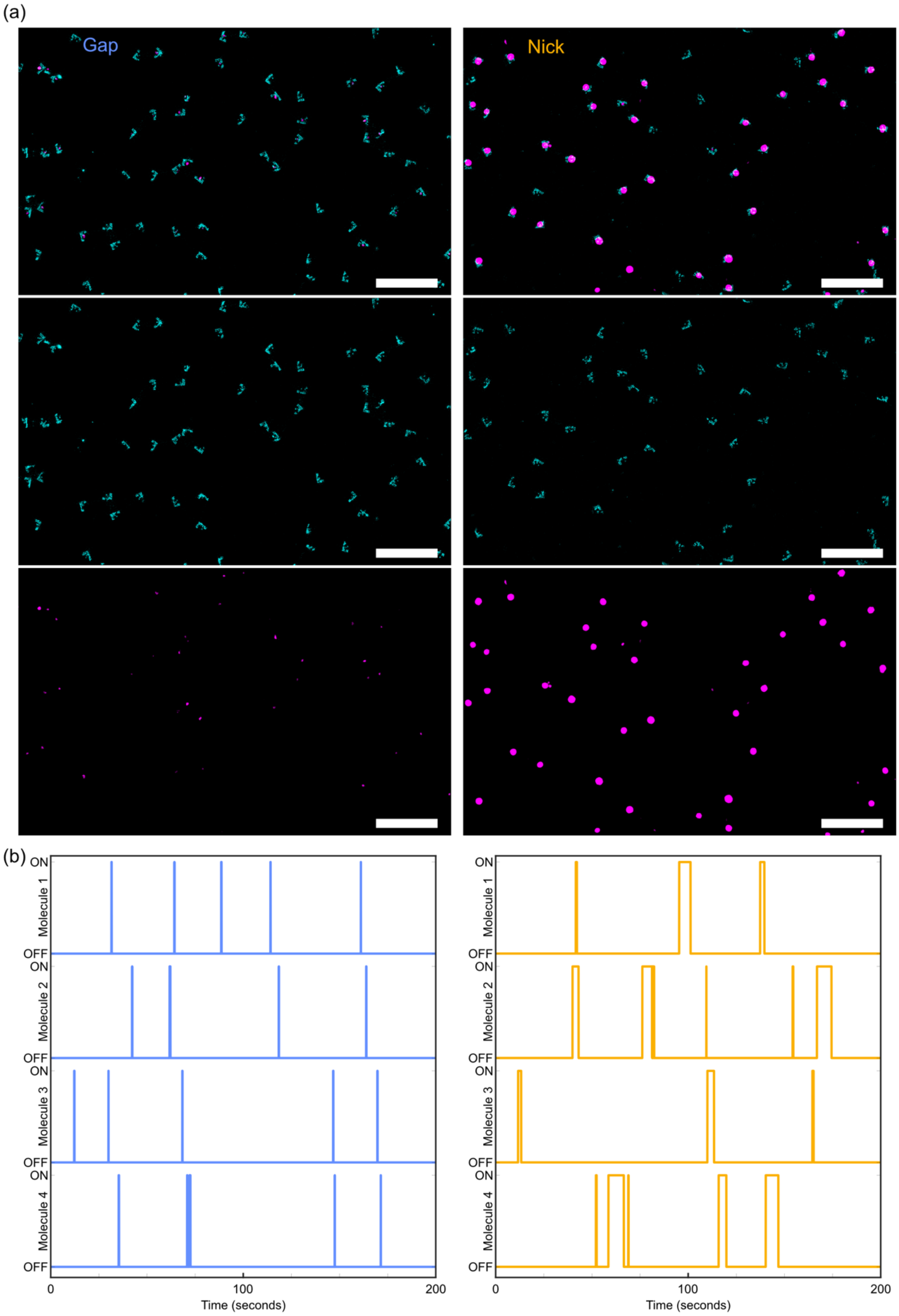


**Supplementary Figure 2:** Overview DNA-PAINT images showing the imager strand binding at the assay site (a) Large field of view showing grid and assay site under gap (left) and nick (right). Combined image (top) was obtained by overlaying the DNA-PAINT grid image (middle) with the assay site image (bottom). (b) Binding time traces for four example assay sites showing short bindings in the case of gap (left) and long bindings in the case of nick (right).

Supplementary Figure 3

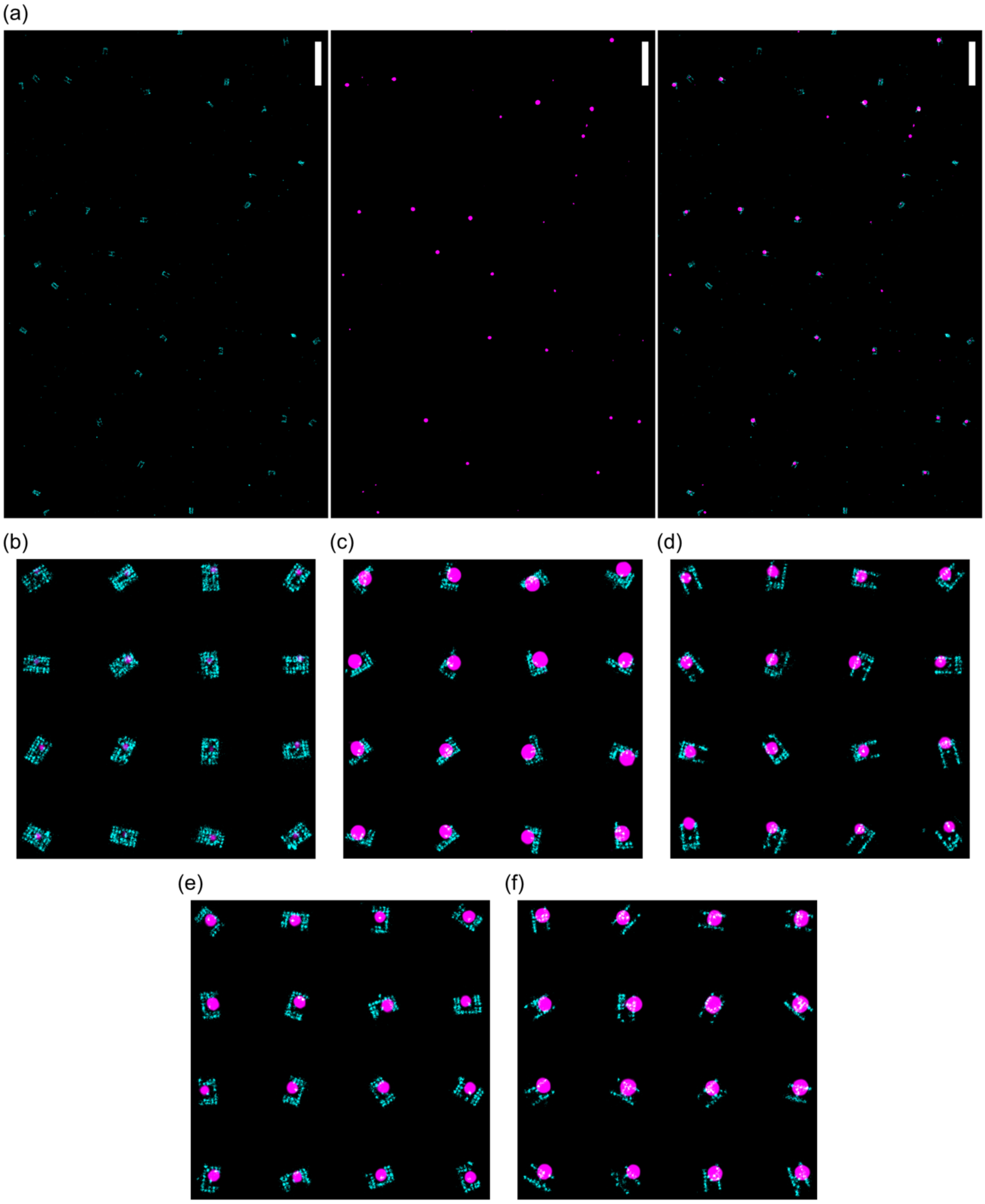


**Supplementary Figure 3:** DNA-PAINT images of simultaneous imaging showing the selected origami structures and corresponding assay site (a) Representative large field of view showing all five grid shapes. As mentioned earlier, we first image the grids with an imager strand carrying ATTO647N (left) following which we image the assay site with an imager strand carrying Cy3B (centre). These two channels are reconstructed individually and then merged (right) for manual picking and further analysis. (Scale = 500nm); Montage of individually picked and aligned origami structures structures showing (b) box-shape for gap, (c) 'L'-shape for stem with A, (d) 'U'-shape for stem with T, (e) 'C'-shape for stem with C, (f) 'H'-shape for stem with G.

Supplementary Figure 4

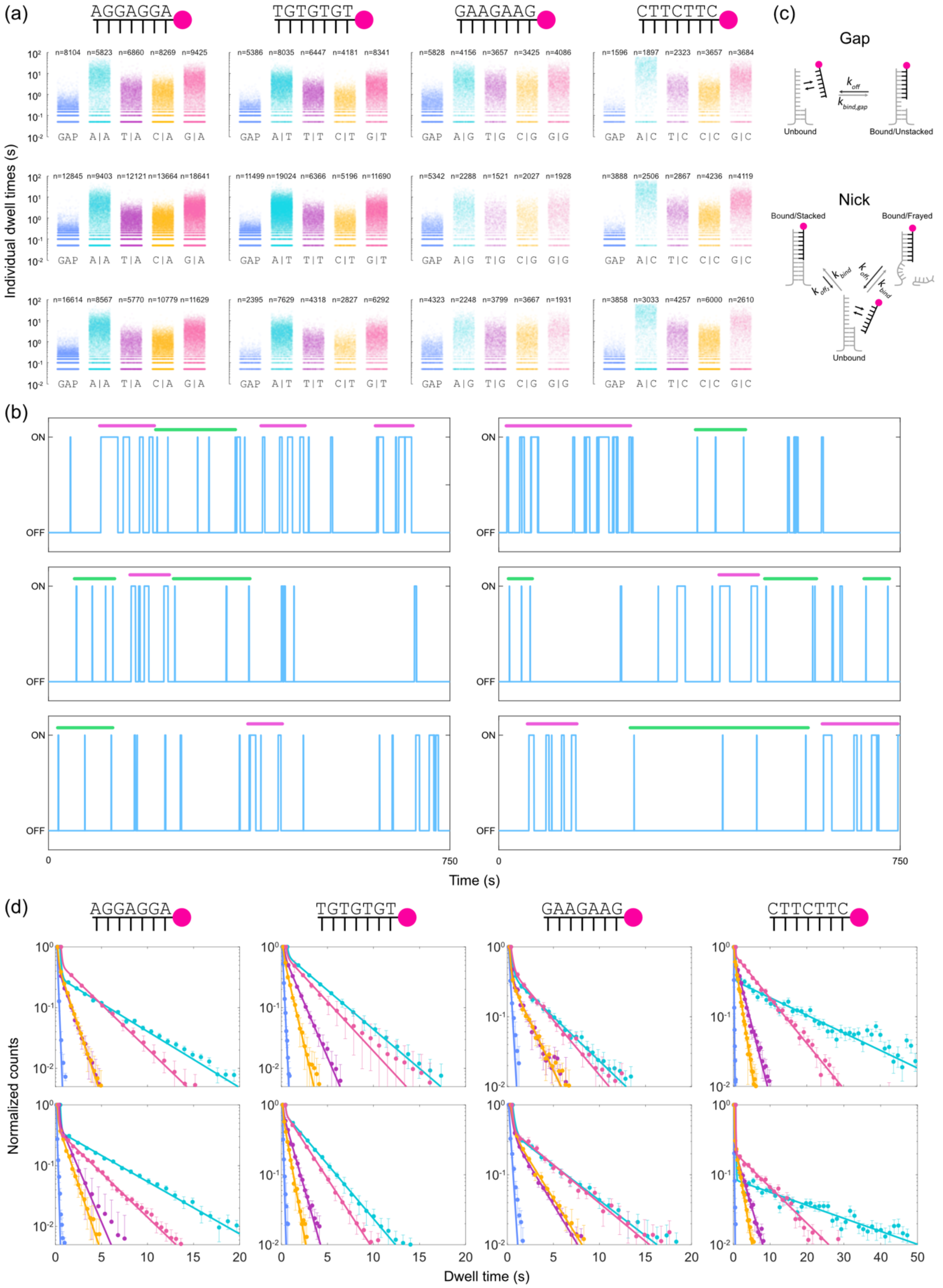


**Supplementary Figure 4:** Dwell times analysis measured under different stacking conditions. (a) Distribution of individual dwell times plotted over a defined randomised spread for each population. Each spot is plotted with a 97% transparency to graphically represent the density spread of the dwell times. These data are related to figure 3a. (b) Representative idealized time traces of imager binding at the assay site showing consecutive short-

and long-lived dwell times. Green line indicates spans of short-lived and magenta line indicates long-lived dwell times. (c) Kinetic model describing the binding configurations of imager binding at the assay site. Imager binding to gap configuration is described by a two-state kinetic model. On the configuration, the imager binding resulted in consecutive short-lived and long lived binding times made us to attribute the short lived population to imager binding while stem frayed. The shorter binding events were spanned for a considerable duration, indicating the frayed stem does not reanneal within a binding event. The imager binding rate would be minimally affected by this. But, the rate of dissociation is dependent on the stem state –frayed or intact–leading to equivalent of the gap and stack interactions, respectively. (d) Replicate-2 and replicate-3 data of figure (3a). Mono- and bi- exponential fits for individual dwell times under gap and different stackings, respectively. Color coding is the same as in (a).

Supplementary Figure 5

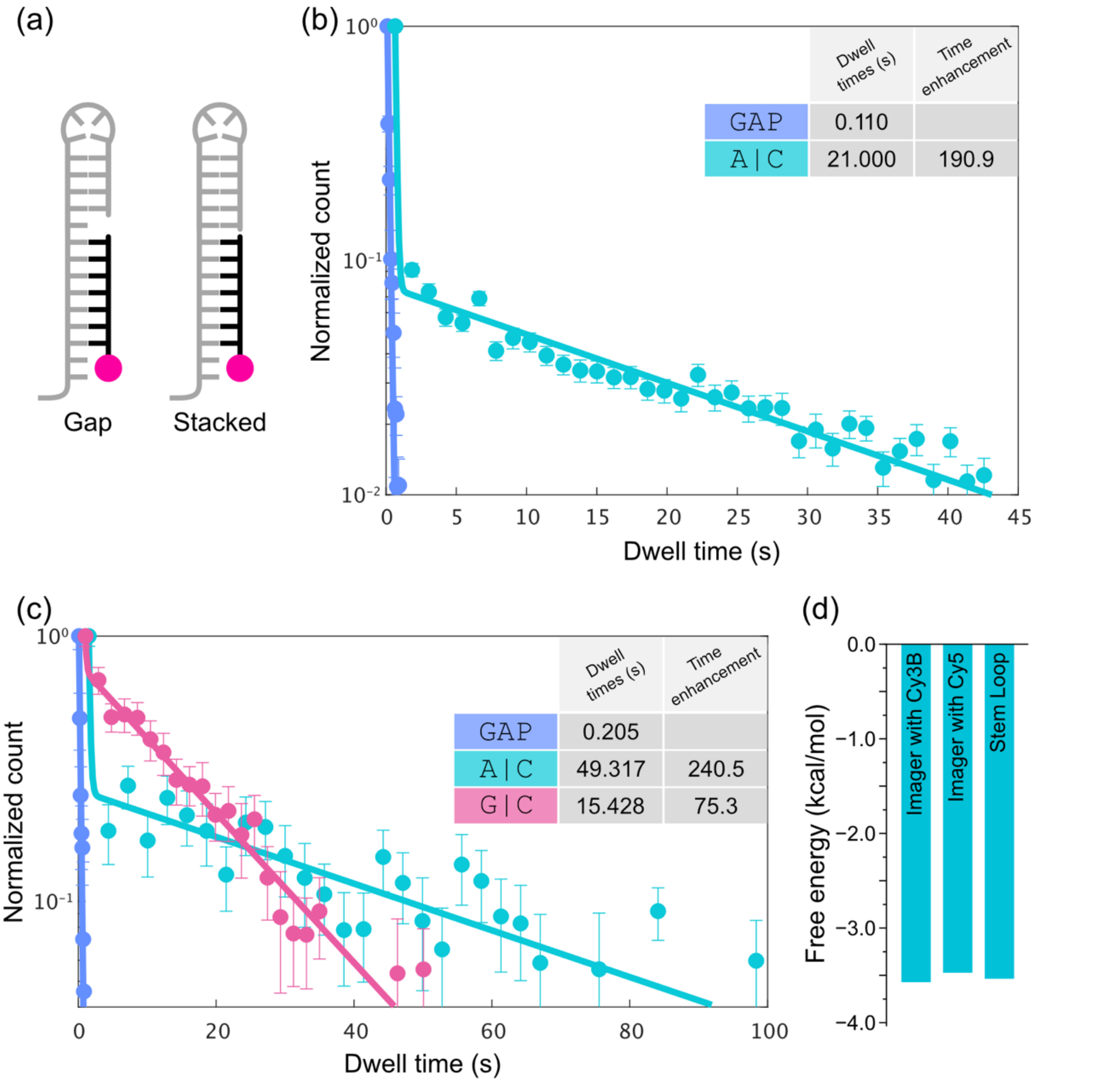


**Supplementary Figure 5:** Orthogonal configurations used to measure A|C stacking interactions. (a) Graphical representation of the orthogonal stem configuration. In this configuration the stem is formed by hairpin formation from the docking strand extension. This configuration carries different stem sequence, providing orthogonal local context in which the stacking energy is measured. (b) Mono- and bi-exponential fits of individual dwell times of the gap and A|C stack, respectively. Dwell times and time enhancements show similar values under this orthogonal condition. (c) Mono- and bi- exponential fits of individual dwell times for the gap and nick configurations, respectively for Cy5-labeled imager. This showed longer discrete binding times as compared to imager containing Cy3B but showed similar time enhancements upon normalizing with the gap average dwell time. (d) Free energy values for A|C stack obtained from measurements with orthogonal configurations. (Compare with Fig 3b)

Supplementary Figure 6

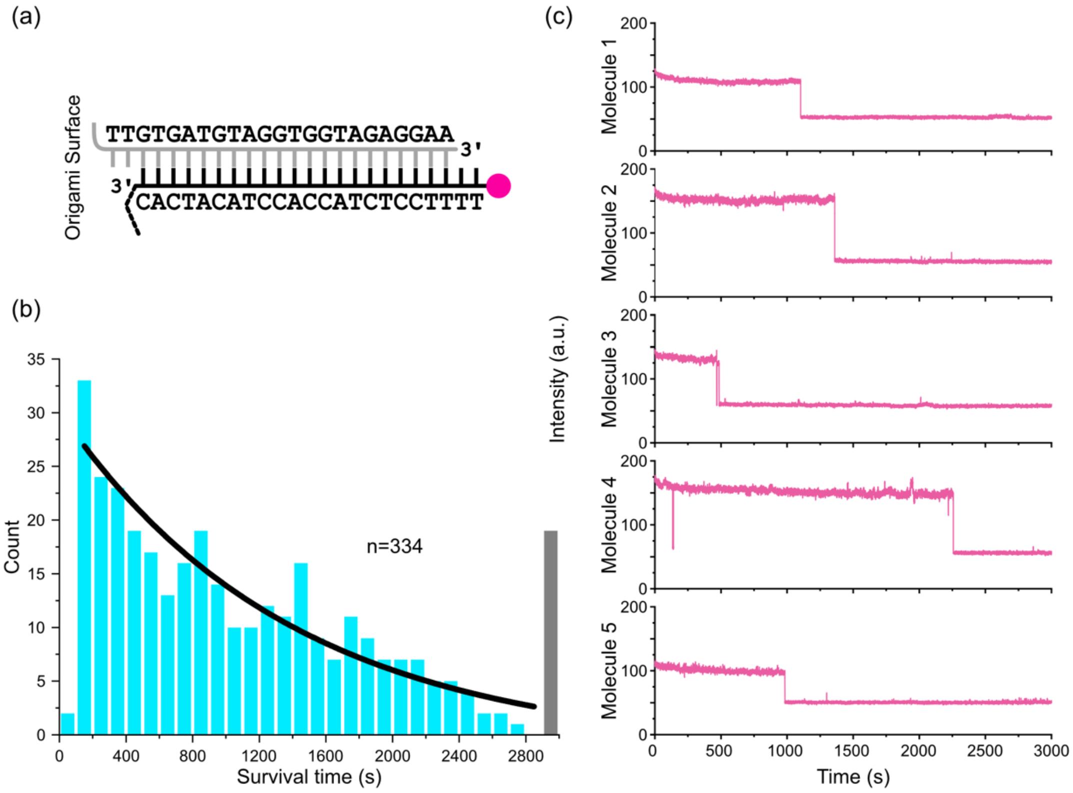


**Supplementary Figure 6:** (a) Graphical representation of the stable duplex used in the photobleaching experiment. (b) Plot showing number of Cy3B fluorophore molecules surviving for a given amount of time (n=313) before photobleaching. Single exponential decay fitting was performed on the histogram plot to obtain the bleaching rate of the fluorophore (0.0007/second). This is substantially lower than the obtained off rates for various stacks, ruling out any requirements of correction for photobleaching in the nick imaging studies. The first bar is ignored for the fitting. The colored bar represent the number of survived (fluorescing until the end of the imaging duration) molecules. (c) Intensity time traces for five individual molecules that showed single step fluorescence disappearance due to photobleaching.

Supplementary Figure 7

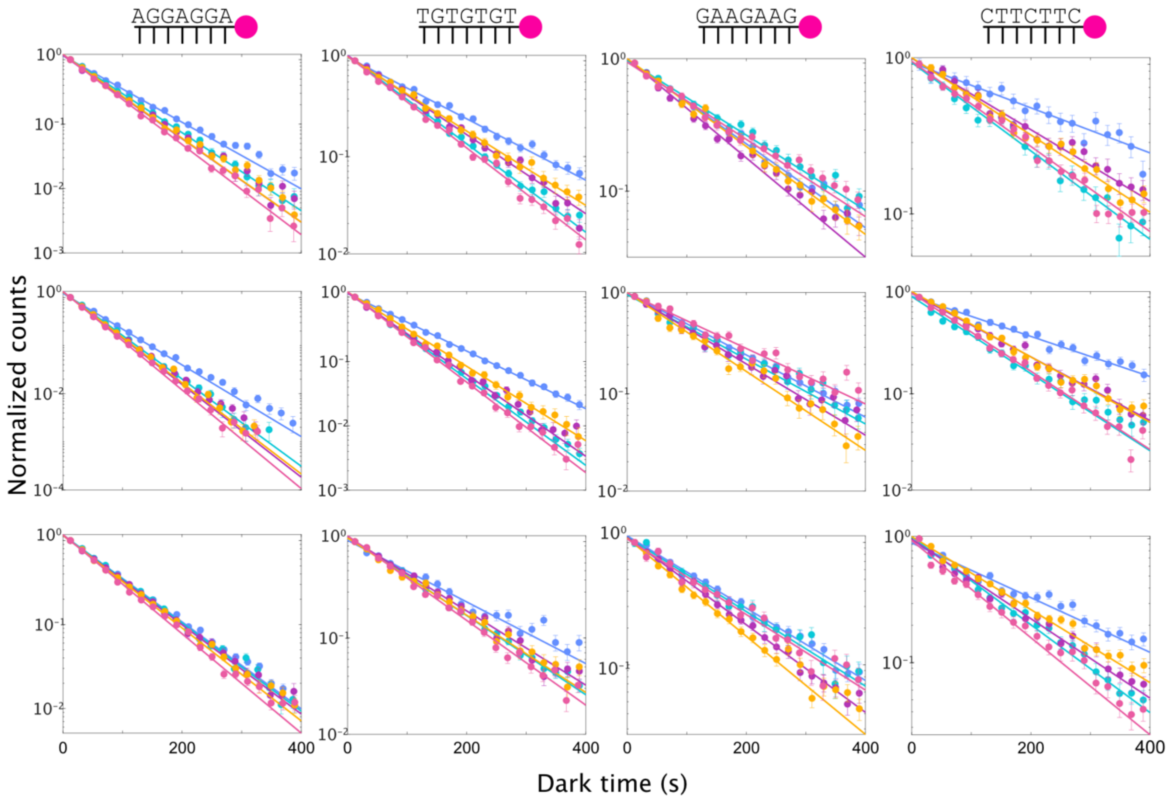


**Supplementary Figure 7:** (a) Histograms of dark time (points) fit with single exponential function (lines) of all four different imager sequences. Each row represents a replicate.

Supplementary Figure 8

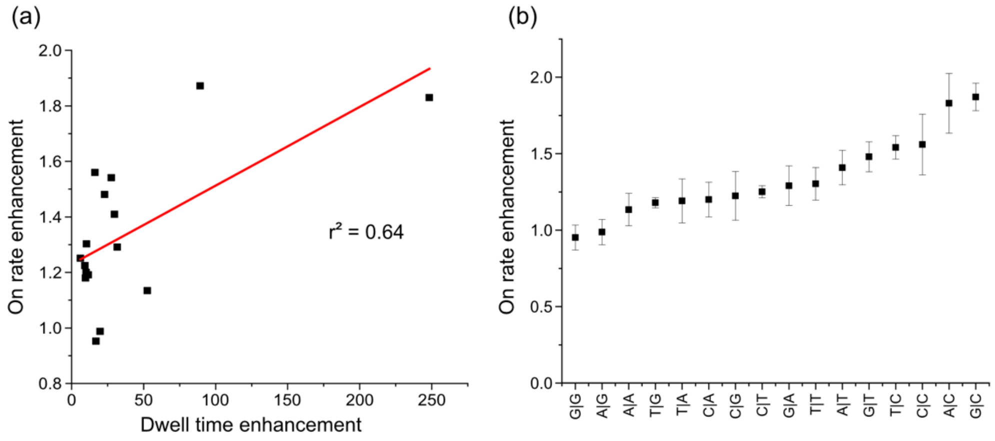


**Supplementary Figure 8:** (a) Plot showing correlation between the dwell time enhancements and the on-rate enhancements. There is a positive correlation between the dwell time enhancements and the on-rate enhancements. ($r^2 = 0.64$) (b) Increase order of on-rate enhancement. Greater on-rate enhancements were clearly seen for stacks with pyrimidine|pyrimidine dinucleotides. Least on-rate enhancements were clearly noted in the case of stack with purine|purine dinucleotides. This would likely arise due to the bulky nature of the purines causing steric hindrance effects on imager binding.

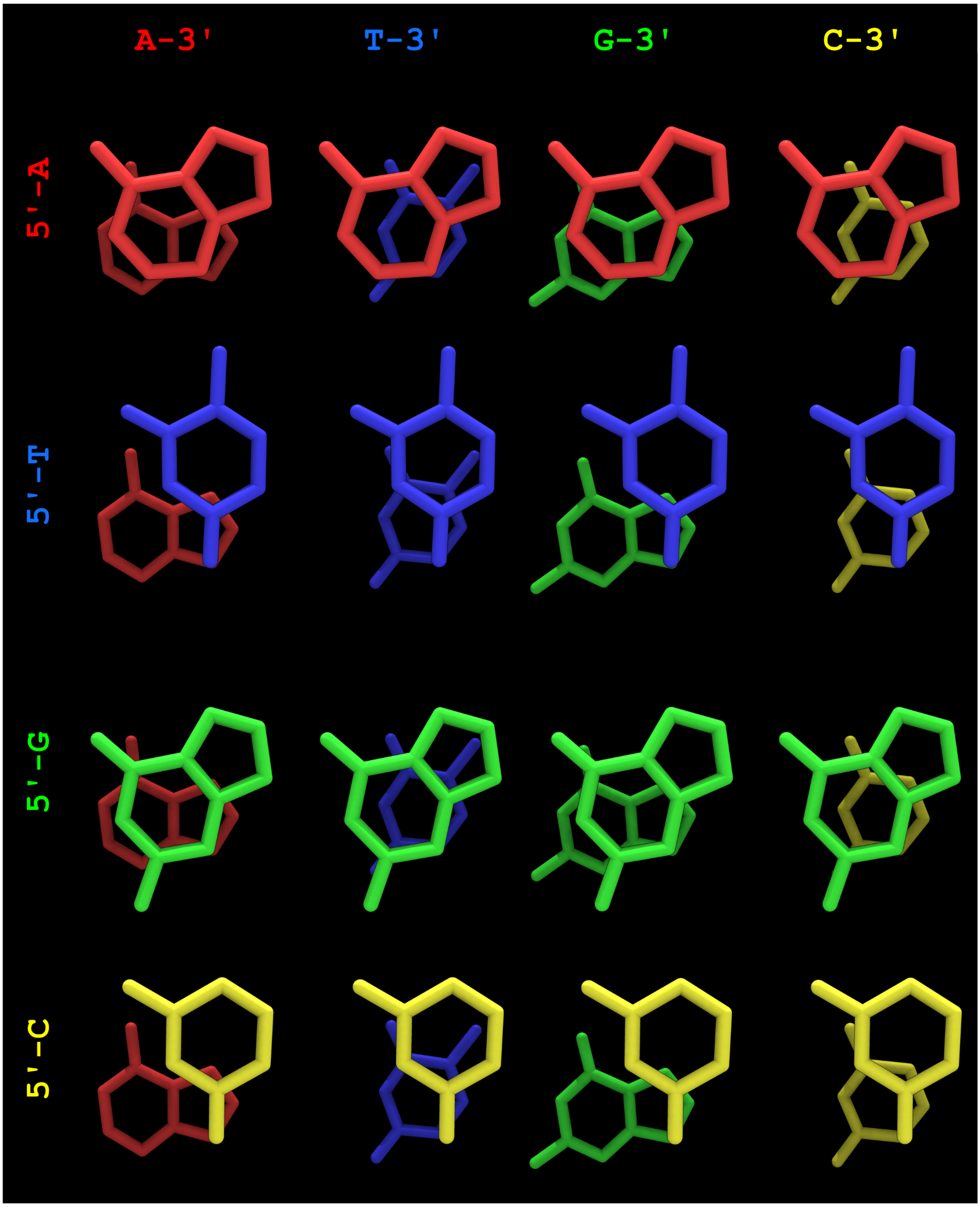


**Supplementary Figure 9:** Top-view of nucleotide positioning from conventional B-form DNA for all 16 combinations. Foreground nucleotide is the nucleotide on the 5' side. The background nucleotide is the nucleotide on the 3' side. These combinations show clear distinction induced by directionality of the sequence. For example, A|C show larger overlap than C|A, in corroboration with around 250 and 10-fold off-rate enhancement, respectively.

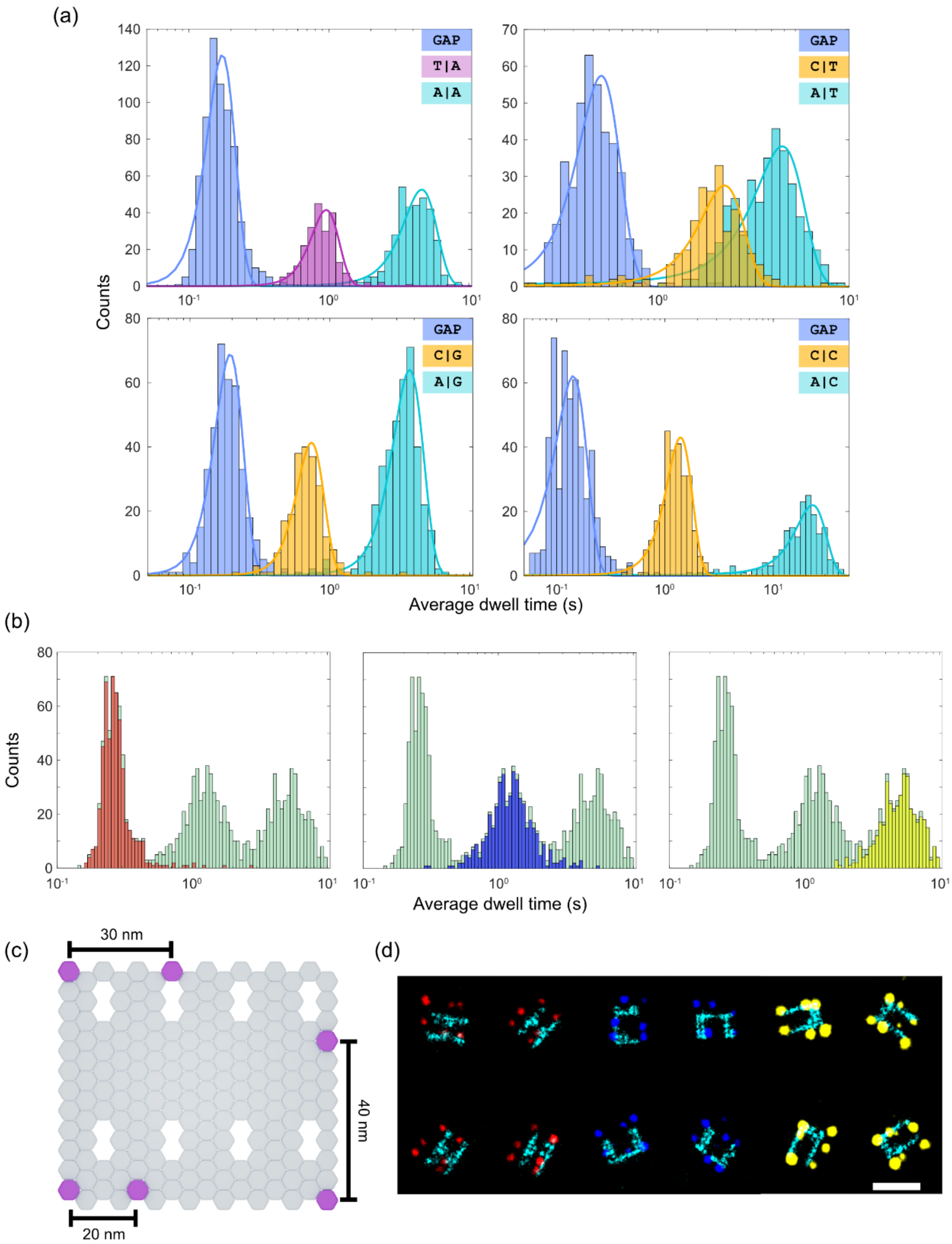


**Supplementary Figure 10:** Stack-PAINT for decoding of several targets with a single imager based on the varying average dwell times of the imager depending on the stacking partner. (a) Histograms showing the distribution of average dwell time for each of the four imagers used in figure 3a. Out of five configurations (a gap and four nicks), we show three potential pairs of stacking configurations that can be used for Stack-PAINT. (b) Manually picked origami based on the ground-truth grid structures show average binding times in three distinguishable peaks. Gap (red), T|A (blue), and A|A (yellow) show distinct average dwell time of the imager on the target. (c) Extensions on origami structures separated by varying distances used to measure the resolution potential of Stack-PAINT. (d) Picked origami structures showing the ground-truth grid structures along with the decoded target spots showing clear distinct 20 nm or farther separated spots with Stack-PAINT.

### Supplementary tables

**Supplementary table 1:** Off rate constants resulted from the exponential fitting for the data sets in figure 3 and supplementary figure 4. Off rate constants, obtained by taking the inverse of dissociation rate ($k_{off,2}$), and dwell time enhancement over gap for each replicate set.

| | Stack | First rate constant ($k_{off}$ or $k_{off,1}$) | Second rate constant ($k_{off,2}$) | Dwell time enhancement | Stack | First rate constant ($k_{off}$ or $k_{off,1}$) | Second rate constant ($k_{off,2}$) | Dwell time enhancement |
|---|---|---|---|---|---|---|---|---|
| Replicate 1 | **GAP** | 6.915 | | | **GAP** | 9.919 | | |
| | **A\|A** | 7.137 | 0.122 | 56.417 | **A\|T** | 11.900 | 0.297 | 33.359 |
| | **T\|A** | 7.165 | 0.594 | 11.629 | **T\|T** | 8.919 | 0.837 | 11.843 |
| | **C\|A** | 7.165 | 0.659 | 10.493 | **C\|T** | 11.920 | 1.431 | 6.929 |
| | **G\|A** | 6.415 | 0.206 | 33.539 | **G\|T** | 11.789 | 0.388 | 25.517 |
| Replicate 2 | **GAP** | 8.866 | | | **GAP** | 7.435 | | |
| | **A\|A** | 8.366 | 0.212 | 41.717 | **A\|T** | 6.935 | 0.296 | 25.057 |
| | **T\|A** | 8.664 | 0.993 | 8.924 | **T\|T** | 6.935 | 0.814 | 9.130 |
| | **C\|A** | 8.366 | 1.056 | 8.393 | **C\|T** | 6.935 | 1.480 | 5.022 |
| | **G\|A** | 8.979 | 0.336 | 26.366 | **G\|T** | 6.935 | 0.376 | 19.758 |
| Replicate 3 | **GAP** | 11.854 | | | **GAP** | 11.064 | | |
| | **A\|A** | 12.064 | 0.198 | 59.83025 | **A\|T** | 12.821 | 0.358 | 30.855 |
| | **T\|A** | 12.069 | 0.816327 | 14.52162 | **T\|T** | 10.064 | 1.061 | 10.425 |
| | **C\|A** | 12.104 | 1.030 | 11.508 | **C\|T** | 12.231 | 1.711 | 6.465 |
| | **G\|A** | 11.354 | 0.335 | 35.284 | **G\|T** | 13.062 | 0.469 | 23.578 |
| Replicate 1 | **GAP** | 4.056 | | | **GAP** | 12.072 | | |
| | **A\|G** | 3.726 | 0.191 | 21.212 | **A\|C** | 12.538 | 0.048 | 247.329 |
| | **T\|G** | 3.056 | 0.358 | 11.321 | **T\|C** | 11.078 | 0.451 | 26.732 |
| | **C\|G** | 3.919 | 0.408 | 9.935 | **C\|C** | 11.072 | 0.736 | 16.382 |
| | **G\|G** | 3.056 | 0.228 | 17.743 | **G\|C** | 12.566 | 0.148 | 81.344 |
| Replicate 2 | **GAP** | 5.333 | | | **GAP** | 13.357 | | |
| | **A\|G** | 4.333 | 0.290 | 18.352 | **A\|C** | 13.779 | 0.057 | 230.371 |
| | **T\|G** | 4.333 | 0.633 | 8.417 | **T\|C** | 15.357 | 0.471 | 28.345 |
| | **C\|G** | 6.753 | 0.632 | 8.433 | **C\|C** | 15.357 | 0.794 | 16.808 |
| | **G\|G** | 4.333 | 0.360 | 14.795 | **G\|C** | 15.349 | 0.149 | 89.595 |
| Replicate 3 | **GAP** | 4.667 | | | **GAP** | 11.482 | | |
| | **A\|G** | 4.244 | 0.233 | 20 | **A\|C** | 13.436 | 0.0429 | 267.486 |
| | **T\|G** | 3.667 | 0.500 | 9.321 | **T\|C** | 10.482 | 0.414 | 27.690 |
| | **C\|G** | 3.667 | 0.486 | 9.593 | **C\|C** | 10.482 | 0.730 | 15.724 |
| | **G\|G** | 6.667 | 0.252 | 18.514 | **G\|C** | 13.340 | 0.118 | 96.760 |

**Supplementary table 2:** Binding rate constants ($k_{bind}$) for all data sets shown in Supplementary figure 7 and enhancement in $k_{bind}$ (i.e., ratio of stack to gap binding rates).

| | Stack | Rate constant | Enhancement in binding rate | Stack | Rate constant | Enhancement in binding rate |
|---|---|---|---|---|---|---|
| Replicate 1 | **GAP** | 0.011 | | **GAP** | 0.007 | |
| | **A\|A** | 0.034 | 0.853 | **A\|T** | 0.010 | 0.694 |
| | **T\|A** | 0.014 | 0.797 | **T\|T** | 0.009 | 0.776 |
| | **C\|A** | 0.014 | 0.793 | **C\|T** | 0.008 | 0.826 |
| | **G\|A** | 0.016 | 0.738 | **G\|T** | 0.011 | 0.667 |
| Replicate 2 | **GAP** | 0.017 | | **GAP** | 0.010 | |
| | **A\|A** | 0.020 | 0.821 | **A\|T** | 0.015 | 0.665 |
| | **T\|A** | 0.022 | 0.772 | **T\|T** | 0.014 | 0.706 |
| | **C\|A** | 0.021 | 0.786 | **C\|T** | 0.013 | 0.775 |
| | **G\|A** | 0.023 | 0.726 | **G\|T** | 0.016 | 0.637 |
| Replicate 3 | **GAP** | 0.011 | | **GAP** | 0.00 | |
| | **A\|A** | 0.012 | 0.985 | **A\|T** | 0.009 | 0.778 |
| | **T\|A** | 0.012 | 0.973 | **T\|T** | 0.008 | 0.830 |
| | **C\|A** | 0.012 | 0.935 | **C\|T** | 0.009 | 0.797 |
| | **G\|A** | 0.013 | 0.875 | **G\|T** | 0.010 | 0.727 |
| Replicate 1 | **GAP** | 0.007 | | **GAP** | 0.003 | |
| | **A\|G** | 0.006 | 1.119 | **A\|C** | 0.006 | 0.510 |
| | **T\|G** | 0.008 | 0.845 | **T\|C** | 0.005 | 0.631 |
| | **C\|G** | 0.007 | 0.954 | **C\|C** | 0.005 | 0.583 |
| | **G\|G** | 0.006 | 1.086 | **G\|C** | 0.006 | 0.529 |
| Replicate 2 | **GAP** | 0.007 | | **GAP** | 0.004 | |
| | **A\|G** | 0.007 | 0.961 | **A\|C** | 0.009 | 0.520 |
| | **T\|G** | 0.008 | 0.873 | **T\|C** | 0.007 | 0.630 |
| | **C\|G** | 0.009 | 0.788 | **C\|C** | 0.007 | 0.613 |
| | **G\|G** | 0.006 | 1.121 | **G\|C** | 0.009 | 0.512 |
| Replicate 3 | **GAP** | 0.006 | | **GAP** | 0.005 | |
| | **A\|G** | 0.007 | 0.970 | **A\|C** | 0.008 | 0.622 |
| | **T\|G** | 0.008 | 0.825 | **T\|C** | 0.007 | 0.688 |
| | **C\|G** | 0.009 | 0.736 | **C\|C** | 0.006 | 0.748 |
| | **G\|G** | 0.007 | 0.956 | **G\|C** | 0.009 | 0.563 |

**Supplementary table 3:** $\Delta G_{stack}$ comparison from different studies. First three columns are free energy of base-pair stacking interactions extracted from unified model for the nearest neighbor parameters[6], gel migrated DNA with nick and gap configurations[7], and blunt end interactions of DNA bundles extrapolated to zero-force and single-pair base-stacks from optical tweezers experiments[8], respectively. We have taken these numbers from reference.[9] Third is the free energy calculation of individual dinucleotide base-stacking interactions from the current study. Fourth column is the average of reverse complements of dinucleotide base-stacks from this study.

| Base-pair | $\Delta G_{,SantaLucia}$ (kcal.mol$^{-1}$) | $\Delta G_{,Nick}$ (kcal.mol$^{-1}$) | $\Delta G_{,Blunt}$ (kcal.mol$^{-1}$) | $\Delta G$ (kcal.mol$^{-1}$) | $\Delta G$ (Average) (kcal.mol$^{-1}$) |
|---|---|---|---|---|---|
| A\|A, T\|T | −1.00 | −1.11 | −1.36 | -2.38, -1.52 | -1.95 |
| AT | −0.88 | −1.34 | −2.35 | -2.17 | -2.17 |
| TA | −0.58 | −0.19 | −1.01 | -1.52 | -1.52 |
| G\|G, C\|C | −1.84 | −1.44 | −1.64 | -1.62, -1.89 | -1.76 |
| GC | −2.24 | −2.17 | −3.42 | -2.99 | -2.99 |
| CG | −2.17 | −0.91 | −2.06 | -1.42 | -1.42 |
| T\|G, C\|A | −1.45 | −0.55 | −0.81 | -1.42, -1.45 | -1.44 |
| A\|G, C\|T | −1.28 | −1.06 | −1.60 | -1.74, -1.19 | -1.46 |
| T\|C, G\|A | −1.30 | −1.43 | −1.39 | -2.19, -2.16 | -2.18 |
| G\|T, A\|C | −1.44 | −1.81 | −2.03 | -2.06, -3.57 | -2.81 |

**Supplementary table 4:** Blank Staples for DNA origami structures

| Staple Name | Staple Sequence | Manufacturer |
|---|---|---|
| **Common Blank Staples** | | |
| 19[32]21[31]BLK | GTCGACTTCGGCCAACGCGCGGGGTTTTTC | Sigma |
| 17[32]19[31]BLK | TGCATCTTTCCCAGTCACGACGGCCTGCAG | Sigma |
| 15[32]17[31]BLK | TAATCAGCGGATTGACCGTAATCGTAACCG | Sigma |
| 13[32]15[31]BLK | AACGCAAAATCGATGAACGGTACCGGTTGA | Sigma |
| 11[32]13[31]BLK | AACAGTTTTGTACCAAAAACATTTTATTTC | Sigma |
| 9[32]11[31]BLK | TTTACCCCAACATGTTTTAAATTTCCATAT | Sigma |
| 7[32]9[31]BLK | TTTAGGACAAATGCTTTAAACAATCAGGTC | Sigma |
| 5[32]7[31]BLK | CATCAAGTAAAACGAACTAACGAGTTGAGA | Sigma |
| 3[32]5[31]BLK | AATACGTTTGAAAGAGGACAGACTGACCTT | Sigma |
| 1[32]3[31]BLK | AGGCTCCAGAGGCTTTGAGGACACGGGTAA | Sigma |
| 23[32]22[48]BLK | CAAATCAAGTTTTTTGGGGTCGAAACGTGGA | Sigma |
| 20[47]18[48]BLK | TTAATGAACTAGAGGATCCCCGGGGGGTAACG | Sigma |
| 16[47]14[48]BLK | ACAAACGGAAAAGCCCCAAAAACACTGGAGCA | Sigma |
| 12[47]10[48]BLK | TAAATCGGGATTCCCAATTCTGCGATATAATG | Sigma |
| 8[47]6[48]BLK | ATCCCCCTATACCACATTCAACTAGAAAAATC | Sigma |
| 4[47]2[48]BLK | GACCAACTAATGCCACTACGAAGGGGGTAGCA | Sigma |
| 2[47]0[48]BLK | ACGGCTACAAAAGGAGCCTTTAATGTGAGAAT | Sigma |
| 21[56]23[63]BLK | AGCTGATTGCCCTTCAGAGTCCACTATTAAAGGGTGCCGT | Sigma |
| 15[64]18[64]BLK | GTATAAGCCAACCCGTCGGATTCTGACGACAGTATCGGCCGCAAGGCG | Sigma |
| 13[64]15[63]BLK | TATATTTTGTCATTGCCTGAGAGTGGAAGATT | Sigma |
| 11[64]13[63]BLK | GATTTAGTCAATAAAGCCTCAGAGAACCCTCA | Sigma |

| | | |
|---|---|---|
| 9[64]11[63]BLK | CGGATTGCAGAGCTTAATTGCTGAAACGAGTA | Sigma |
| 7[56]9[63]BLK | ATGCAGATACATAACGGGAATCGTCATAAATAAAGCAAAG | Sigma |
| 1[64]4[64]BLK | TTTATCAGGACAGCATCGGAACGACACCAACCTAAAACGAGGTCAATC | Sigma |
| 0[79]1[63]BLK | ACAACTTTCAACAGTTTCAGCGGATGTATCGG | Sigma |
| 23[64]22[80]BLK | AAAGCACTAAATCGGAACCCTAATCCAGTT | Sigma |
| 20[79]18[80]BLK | TTCCAGTCGTAATCATGGTCATAAAAGGGG | Sigma |
| 16[79]14[80]BLK | GCGAGTAAAAATATTTAAATTGTTACAAAG | Sigma |
| 12[79]10[80]BLK | AAATTAAGTTGACCATTAGATACTTTTGCG | Sigma |
| 8[79]6[80]BLK | AATACTGCCCAAAAGGAATTACGTGGCTCA | Sigma |
| 4[79]2[80]BLK | GCGCAGACAAGAGGCAAAAGAATCCCTCAG | Sigma |
| 2[79]0[80]BLK | CAGCGAAACTTGCTTTCGAGGTGTTGCTAA | Sigma |
| 21[96]23[95]BLK | AGCAAGCGTAGGGTTGAGTGTTGTAGGGAGCC | Sigma |
| 19[96]21[95]BLK | CTGTGTGATTGCGTTGCGCTCACTAGAGTTGC | Sigma |
| 17[96]19[95]BLK | GCTTTCCGATTACGCCAGCTGGCGGCTGTTTC | Sigma |
| 15[96]17[95]BLK | ATATTTTGGCTTTCATCAACATTATCCAGCCA | Sigma |
| 13[96]15[95]BLK | TAGGTAAACTATTTTTGAGAGATCAAACGTTA | Sigma |
| 11[96]13[95]BLK | AATGGTCAACAGGCAAGGCAAAGAGTAATGTG | Sigma |
| 9[96]11[95]BLK | CGAAAGACTTTGATAAGAGGTCATATTTCGCA | Sigma |
| 7[96]9[95]BLK | TAAGAGCAAATGTTTAGACTGGATAGGAAGCC | Sigma |
| 5[96]7[95]BLK | TCATTCAGATGCGATTTTAAGAACAGGCATAG | Sigma |
| 3[96]5[95]BLK | ACACTCATCCATGTTACTTAGCCGAAAGCTGC | Sigma |
| 1[96]3[95]BLK | AAACAGCTTTTTGCGGGATCGTCAACACTAAA | Sigma |
| 23[96]22[112]BLK | CCCGATTTAGAGCTTGACGGGGAAAAAGAATA | Sigma |
| 20[111]18[112]BLK | CACATTAAAATTGTTATCCGCTCATGCGGGCC | Sigma |
| 16[111]14[112]BLK | TGTAGCCATTAAAATTCGCATTAAATGCCGGA | Sigma |
| 14[111]12[112]BLK | GAGGGTAGGATTCAAAAGGGTGAGACATCCAA | Sigma |
| 12[111]10[112]BLK | TAAATCATATAACCTGTTTAGCTAACCTTTAA | Sigma |
| 8[111]6[112]BLK | AATAGTAAACACTATCATAACCCTCATTGTGA | Sigma |
| 4[111]2[112]BLK | GACCTGCTCTTTGACCCCCAGCGAGGGAGTTA | Sigma |
| 2[111]0[112]BLK | AAGGCCGCTGATACCGATAGTTGCGACGTTAG | Sigma |
| 15[128]18[128]BLK | TAAATCAAAATAATTCGCGTCTCGGAAACCAGGCAAAGGGAAGG | Sigma |
| 13[128]15[127]BLK | GAGACAGCTAGCTGATAAATTAATTTTTGT | Sigma |
| 11[128]13[127]BLK | TTTGGGGATAGTAGTAGCATTAAAAGGCCG | Sigma |
| 9[128]11[127]BLK | GCTTCAATCAGGATTAGAGAGTTATTTTCA | Sigma |
| 7[120]9[127]BLK | CGTTTACCAGACGACAAAGAAGTTTTGCCATAATTCGA | Sigma |
| 1[128]4[128]BLK | TGACAACTCGCTGAGGCTTGCATTATACCAAGCGCGATGATAAA | Sigma |
| 0[143]1[127]BLK | TCTAAAGTTTTGTCGTCTTTCCAGCCGACAA | Sigma |
| 21[160]22[144]BLK | TCAATATCGAACCTCAAATATCAATTCCGAAA | Sigma |
| 17[160]18[144]BLK | AGAAAACAAAGAAGATGATGAAACAGGCTGCG | Sigma |
| 13[160]14[144]BLK | GTAATAAGTTAGGCAGAGGCATTTATGATATT | Sigma |
| 9[160]10[144]BLK | AGAGAGAAAAAAATGAAAATAGCAAGCAAACT | Sigma |
| 5[160]6[144]BLK | GCAAGGCCTCACCAGTAGCACCATGGGCTTGA | Sigma |
| 1[160]2[144]BLK | TTAGGATTGGCTGAGACTCCTCAATAACCGAT | Sigma |
| 0[175]0[144]BLK | TCCACAGACAGCCCTCATAGTTAGCGTAACGA | Sigma |

| | | |
|---|---|---|
| 23[128]23[159]BLK | AACGTGGCGAGAAAGGAAGGGAAACCAGTAA | Sigma |
| 22[143]21[159]BLK | TCGGCAAATCCTGTTTGATGGTGGACCCTCAA | Sigma |
| 20[143]19[159]BLK | AAGCCTGGTACGAGCCGGAAGCATAGATGATG | Sigma |
| 18[143]17[159]BLK | CAACTGTTGCGCCATTCGCCATTCAAACATCA | Sigma |
| 16[143]15[159]BLK | GCCATCAAGCTCATTTTTTAACCACAAATCCA | Sigma |
| 14[143]13[159]BLK | CAACCGTTTCAAATCACCATCAATTCGAGCCA | Sigma |
| 10[143]9[159]BLK | CCAACAGGAGCGAACCAGACCGGAGCCTTTAC | Sigma |
| 8[143]7[159]BLK | CTTTTGCAGATAAAAACCAAAATAAAGACTCC | Sigma |
| 6[143]5[159]BLK | GATGGTTTGAACGAGTAGTAAATTTACCATTA | Sigma |
| 4[143]3[159]BLK | TCATCGCCAACAAAGTACAACGGACGCCAGCA | Sigma |
| 2[143]1[159]BLK | ATATTCGGAACCATCGCCCACGCAGAGAAGGA | Sigma |
| 23[160]22[176]BLK | TAAAAGGGACATTCTGGCCAACAAAGCATC | Sigma |
| 20[175]18[176]BLK | ATTATCATTCAATATAATCCTGACAATTAC | Sigma |
| 16[175]14[176]BLK | TATAACTAACAAAGAACGCGAGAACGCCAA | Sigma |
| 14[175]12[176]BLK | CATGTAATAGAATATAAAGTACCAAGCCGT | Sigma |
| 12[175]10[176]BLK | TTTTATTTAAGCAAATCAGATATTTTTTGT | Sigma |
| 8[175]6[176]BLK | ATACCCAACAGTATGTTAGCAAATTAGAGC | Sigma |
| 4[175]2[176]BLK | CACCAGAAAGGTTGAGGCAGGTCATGAAAG | Sigma |
| 2[175]0[176]BLK | TATTAAGAAGCGGGGTTTTGCTCGTAGCAT | Sigma |
| 21[184]23[191]BLK | TCAACAGTTGAAAGGAGCAAATGAAAAATCTAGAGATAGA | Sigma |
| 15[192]18[192]BLK | TCAAATATAACCTCCGGCTTAGGTAACAATTTCATTTGAAGGCGAATT | Sigma |
| 13[192]15[191]BLK | GTAAAGTAATCGCCATATTTAACAAAACTTTT | Sigma |
| 11[192]13[191]BLK | TATCCGGTCTCATCGAGAACAAGCGACAAAAG | Sigma |
| 9[192]11[191]BLK | TTAGACGGCCAAATAAGAAACGATAGAAGGCT | Sigma |
| 7[184]9[191]BLK | CGTAGAAAATACATACCGAGGAAACGCAATAAGAAGCGCA | Sigma |
| 1[192]4[192]BLK | GCGGATAACCTATTATTCTGAAACAGACGATTGGCCTTGAAGAGCCAC | Sigma |
| 0[207]1[191]BLK | TCACCAGTACAAACTACAACGCCTAGTACCAG | Sigma |
| 23[192]22[208]BLK | ACCCTTCTGACCTGAAAGCGTAAGACGCTGAG | Sigma |
| 20[207]18[208]BLK | GCGGAACATCTGAATAATGGAAGGTACAAAAT | Sigma |
| 16[207]14[208]BLK | ACCTTTTTATTTTAGTTAATTTCATAGGGCTT | Sigma |
| 14[207]12[208]BLK | AATTGAGAATTCTGTCCAGACGACTAAACCAA | Sigma |
| 12[207]10[208]BLK | GTACCGCAATTCTAAGAACGCGAGTATTATTT | Sigma |
| 8[207]6[208]BLK | AAGGAAACATAAAGGTGGCAACATTATCACCG | Sigma |
| 4[207]2[208]BLK | CCACCCTCTATTCACAAACAAATACCTGCCTA | Sigma |
| 2[207]0[208]BLK | TTTCGGAAGTGCCGTCGAGAGGGTGAGTTTCG | Sigma |
| 21[224]23[223]BLK | CTTTAGGGCCTGCAACAGTGCCAATACGTG | Sigma |
| 19[224]21[223]BLK | CTACCATAGTTTGAGTAACATTTAAAATAT | Sigma |
| 17[224]19[223]BLK | CATAAATCTTTGAATACCAAGTGTTAGAAC | Sigma |
| 15[224]17[223]BLK | CCTAAATCAAAATCATAGGTCTAAACAGTA | Sigma |
| 13[224]15[223]BLK | ACAACATGCCAACGCTCAACAGTCTTCTGA | Sigma |
| 11[224]13[223]BLK | GCGAACCTCCAAGAACGGGTATGACAATAA | Sigma |
| 9[224]11[223]BLK | AAAGTCACAAAATAAACAGCCAGCGTTTTA | Sigma |
| 7[224]9[223]BLK | AACGCAAAGATAGCCGAACAAACCCTGAAC | Sigma |
| 5[224]7[223]BLK | TCAAGTTTCATTAAAGGTGAATATAAAAGA | Sigma |

| | | |
|---|---|---|
| 3[224]5[223]BLK | TTAAAGCCAGAGCCGCCACCCTCGACAGAA | Sigma |
| 1[224]3[223]BLK | GTATAGCAAACAGTTAATGCCCAATCCTCA | Sigma |
| 0[239]1[223]BLK | AGGAACCCATGTACCGTAACACTTGATATAA | Sigma |
| 23[224]22[240]BLK | GCACAGACAATATTTTTGAATGGGGTCAGTA | Sigma |
| 20[239]18[240]BLK | ATTTTAAAATCAAAATTATTTGCACGGATTCG | Sigma |
| 16[239]14[240]BLK | GAATTTATTTAATGGTTTGAAATATTCTTACC | Sigma |
| 12[239]10[240]BLK | CTTATCATTCCCGACTTGCGGGAGCCTAATTT | Sigma |
| 8[239]6[240]BLK | AAGTAAGCAGACACCACGGAATAATATTGACG | Sigma |
| 4[239]2[240]BLK | GCCTCCCTCAGAATGGAAAGCGCAGTAACAGT | Sigma |
| 2[239]0[240]BLK | GCCCGTATCCGGAATAGGTGTATCAGCCCAAT | Sigma |
| 21[248]23[255]BLK | AGATTAGAGCCGTCAAAAAACAGAGGTGAGGCCTATTAGT | Sigma |
| 15[256]18[256]BLK | GTGATAAAAAGACGCTGAGAAGAGATAACCTTGCTTCTGTTCGGGAGA | Sigma |
| 13[256]15[255]BLK | GTTTATCAATATGCGTTATACAAACCGACCGT | Sigma |
| 11[256]13[255]BLK | GCCTTAAACCAATCAATAATCGGCACGCGCCT | Sigma |
| 9[256]11[255]BLK | GAGAGATAGAGCGTCTTTCCAGAGGTTTTGAA | Sigma |
| 7[248]9[255]BLK | GTTTATTTTGTCACAATCTTACCGAAGCCCTTTAATATCA | Sigma |
| 1[256]4[256]BLK | CAGGAGGTGGGGTCAGTGCCTTGAGTCTCTGAATTTACCGGGAACCAG | Sigma |
| 0[271]1[255]BLK | CCACCCTCATTTTCAGGGATAGCAACCGTACT | Sigma |
| 23[256]22[272]BLK | CTTTAATGCGCGAACTGATAGCCCCACCAG | Sigma |
| 20[271]18[272]BLK | CTCGTATTAGAAATTGCGTAGATACAGTAC | Sigma |
| 16[271]14[272]BLK | CTTAGATTTAAGGCGTTAAATAAAGCCTGT | Sigma |
| 12[271]10[272]BLK | TGTAGAAATCAAGATTAGTTGCTCTTACCA | Sigma |
| 8[271]6[272]BLK | AATAGCTATCAATAGAAAATTCAACATTCA | Sigma |
| 4[271]2[272]BLK | AAATCACCTTCCAGTAAGCGTCAGTAATAA | Sigma |
| **Blank Staples specific for 'C'-shape** | | |
| 18[111]16[112]BLK | TCTTCGCTGCACCGCTTCTGGTGCGGCCTTCC | Sigma |
| 10[111]8[112]BLK | TTGCTCCTTTCAAATATCGCGTTTGAGGGGGT | Sigma |
| 6[111]4[112]BLK | ATTACCTTTGAATAAGGCTTGCCCAAATCCGC | Sigma |
| 15[160]16[144]BLK | ATCGCAAGTATGTAAATGCTGATGATAGGAAC | Sigma |
| 7[160]8[144]BLK | TTATTACGAAGAACTGGCATGATTGCGAGAGG | Sigma |
| 3[160]4[144]BLK | TTGACAGGCCACCACCAGAGCCGCGATTTGTA | Sigma |
| 18[175]16[176]BLK | CTGAGCAAAAATTAATTACATTTTGGGTTA | Sigma |
| 10[175]8[176]BLK | TTAACGTCTAACATAAAAACAGGTAACGGA | Sigma |
| 6[175]4[176]BLK | CAGCAAAAGGAAACGTCACCAATGAGCCGC | Sigma |
| 18[207]16[208]BLK | CGCGCAGATTACCTTTTTTAATGGGAGAGACT | Sigma |
| 10[207]8[208]BLK | ATCCCAATGAGAATTAACTGAACAGTTACCAG | Sigma |
| 6[207]4[208]BLK | TCACCGACGCACCGTAATCAGTAGCAGAACCG | Sigma |
| 21[32]23[31]BLK | TTTTCACTCAAAGGGCGAAAAACCATCACC | Sigma |
| 0[47]1[31]BLLK | AGAAAGGAACAACTAAAGGAATTCAAAAAAA | Sigma |
| 0[111]1[95]BLK | TAAATGAATTTTCTGTATGGGATTAATTTCTT | Sigma |
| 21[128]23[127]BLK | CCCAGCAGGCGAAAAATCCCTTATAAATCAAGCCGGCG | Sigma |
| 2[271]0[272]BLK | GTTTTAACTTAGTACCGCCACCCAGAGCCA | Sigma |
| **Blank Staples specific for 'U'-shape** | | |
| 18[111]16[112]BLK | TCTTCGCTGCACCGCTTCTGGTGCGGCCTTCC | Sigma |

| | | |
|---|---|---|
| 10[111]8[112]BLK | TTGCTCCTTTCAAATATCGCGTTTGAGGGGGT | Sigma |
| 15[160]16[144]BLK | ATCGCAAGTATGTAAATGCTGATGATAGGAAC | Sigma |
| 7[160]8[144]BLK | TTATTACGAAGAACTGGCATGATTGCGAGAGG | Sigma |
| 18[175]16[176]BLK | CTGAGCAAAAATTAATTACATTTTGGGTTA | Sigma |
| 8[175]6[176]BLK | TTAACGTCTAACATAAAAACAGGTAACGGA | Sigma |
| 18[207]16[208]BLK | CGCGCAGATTACCTTTTTTAATGGGAGAGACT | Sigma |
| 10[207]8[208]BLK | ATCCCAATGAGAATTAACTGAACAGTTACCAG | Sigma |
| 18[239]16[240]BLK | CCTGATTGCAATATATGTGAGTGATCAATAGT | Sigma |
| 14[239]12[240]BLK | AGTATAAAGTTCAGCTAATGCAGATGTCTTTC | Sigma |
| 10[239]8[240]BLK | GCCAGTTAGAGGGTAATTGAGCGCTTTAAGAA | Sigma |
| 18[271]16[272]BLK | CTTTTACAAAATCGTCGCTATTAGCGATAG | Sigma |
| 14[271]12[272]BLK | TTAGTATCACAATAGATAAGTCCACGAGCA | Sigma |
| 10[271]8[272]BLK | ACGCTAACACCCACAAGAATTGAAAATAGC | Sigma |
| 21[32]23[31]BLK | TTTTCACTCAAAGGGCGAAAAACCATCACC | Sigma |
| 0[47]1[31]BLLK | AGAAAGGAACAACTAAAGGAATTCAAAAAAA | Sigma |
| 0[111]1[95]BLK | TAAATGAATTTTCTGTATGGGATTAATTTCTT | Sigma |
| 21[128]23[127]BLK | CCCAGCAGGCGAAAAATCCCTTATAAATCAAGCCGGCG | Sigma |
| 2[271]0[272]BLK | GTTTTAACTTAGTACCGCCACCCAGAGCCA | Sigma |
| 18[271]16[272]BLK | CTTTTACAAAATCGTCGCTATTAGCGATAG | Sigma |
| **Blank Staples specific for 'H'-shape** | | |
| 18[47]16[48]BLK | CCAGGGTTGCCAGTTTGAGGGGACCCGTGGGA | Sigma |
| 14[47]12[48]BLK | AACAAGAGGGATAAAAATTTTTAGCATAAAGC | Sigma |
| 10[47]8[48]BLK | CTGTAGCTTGACTATTATAGTCAGTTCATTGA | Sigma |
| 18[79]16[80]BLK | GATGTGCTTCAGGAAGATCGCACAATGTGA | Sigma |
| 14[79]12[80]BLK | GCTATCAGAAATGCAATGCCTGAATTAGCA | Sigma |
| 10[79]8[80]BLK | GATGGCTTATCAAAAAGATTAAGAGCGTCC | Sigma |
| 18[111]16[112]BLK | TCTTCGCTGCACCGCTTCTGGTGCGGCCTTCC | Sigma |
| 10[111]8[112]BLK | TTGCTCCTTTCAAATATCGCGTTTGAGGGGGT | Sigma |
| 18[207]16[208]BLK | CGCGCAGATTACCTTTTTTAATGGGAGAGACT | Sigma |
| 10[207]8[208]BLK | ATCCCAATGAGAATTAACTGAACAGTTACCAG | Sigma |
| 18[239]16[240]BLK | CCTGATTGCAATATATGTGAGTGATCAATAGT | Sigma |
| 14[239]12[240]BLK | AGTATAAAGTTCAGCTAATGCAGATGTCTTTC | Sigma |
| 10[239]8[240]BLK | GCCAGTTAGAGGGTAATTGAGCGCTTTAAGAA | Sigma |
| 18[271]16[272]BLK | CTTTTACAAAATCGTCGCTATTAGCGATAG | Sigma |
| 14[271]12[272]BLK | TTAGTATCACAATAGATAAGTCCACGAGCA | Sigma |
| 10[271]8[272]BLK | ACGCTAACACCCACAAGAATTGAAAATAGC | Sigma |
| 21[32]23[31]BLK | TTTTCACTCAAAGGGCGAAAAACCATCACC | Sigma |
| 0[47]1[31]BLLK | AGAAAGGAACAACTAAAGGAATTCAAAAAAA | Sigma |
| 0[111]1[95]BLK | TAAATGAATTTTCTGTATGGGATTAATTTCTT | Sigma |
| 21[128]23[127]BLK | CCCAGCAGGCGAAAAATCCCTTATAAATCAAGCCGGCG | Sigma |
| 2[271]0[272]BLK | GTTTTAACTTAGTACCGCCACCCAGAGCCA | Sigma |
| 18[271]16[272]BLK | CTTTTACAAAATCGTCGCTATTAGCGATAG | Sigma |
| **Blank Staples specific for 'L'-shape** | | |
| 10[111]8[112]BLK | TTGCTCCTTTCAAATATCGCGTTTGAGGGGGT | Sigma |

| | | |
|---|---|---|
| 6[111]4[112]BLK | ATTACCTTTGAATAAGGCTTGCCCAAATCCGC | Sigma |
| 7[160]8[144]BLK | TTATTACGAAGAACTGGCATGATTGCGAGAGG | Sigma |
| 3[160]4[144]BLK | TTGACAGGCCACCACCAGAGCCGCGATTTGTA | Sigma |
| 10[175]8[176]BLK | TTAACGTCTAACATAAAAACAGGTAACGGA | Sigma |
| 6[175]4[176]BLK | CAGCAAAAGGAAACGTCACCAATGAGCCGC | Sigma |
| 10[207]8[208]BLK | ATCCCAATGAGAATTAACTGAACAGTTACCAG | Sigma |
| 6[207]4[208]BLK | TCACCGACGCACCGTAATCAGTAGCAGAACCG | Sigma |
| 14[239]12[240]BLK | AGTATAAAGTTCAGCTAATGCAGATGTCTTTC | Sigma |
| 10[239]8[240]BLK | GCCAGTTAGAGGGTAATTGAGCGCTTTAAGAA | Sigma |
| 6[239]4[240]BLK | GAAATTATTGCCTTTAGCGTCAGACCGGAACC | Sigma |
| 14[271]12[272]BLK | TTAGTATCACAATAGATAAGTCCACGAGCA | Sigma |
| 10[271]8[272]BLK | ACGCTAACACCCACAAGAATTGAAAATAGC | Sigma |
| 6[271]4[272]BLK | ACCGATTGTCGGCATTTTCGGTCATAATCA | Sigma |
| 21[32]23[31]BLK | TTTTCACTCAAAGGGCGAAAAACCATCACC | Sigma |
| 0[47]1[31]BLLK | AGAAAGGAACAACTAAAGGAATTCAAAAAAA | Sigma |
| 0[111]1[95]BLK | TAAATGAATTTTCTGTATGGGATTAATTTCTT | Sigma |
| 21[128]23[127]BLK | CCCAGCAGGCGAAAAATCCCTTATAAATCAAGCCGGCG | Sigma |
| 2[271]0[272]BLK | GTTTTAACTTAGTACCGCCACCCAGAGCCA | Sigma |
| **Blank Staples specific for Barcoded Origami** | | |
| 11[160]12[144]BLK | CCAATAGCTCATCGTAGGAATCATGGCATCAA | Sigma |
| 12[143]11[159]BLK | TTCTACTACGCGAGCTGAAAAGGTTACCGCGC | Sigma |

**Supplementary table 5:** Staples with extensions for grid structures

| Staple Name | Staple Sequence | Manufacturer |
|---|---|---|
| **Staples with R1×5 extensions for 'Box'-shaped grid** | | |
| 22[47]20[48]R1X5 | CTCCAACGCAGTGAGACGGGCAACCAGCTGCA TT TCCTCCTCCTCCTCCTCCT | Sigma |
| 18[47]16[48]R1X5 | CCAGGGTTGCCAGTTTGAGGGGACCCGTGGGA TT TCCTCCTCCTCCTCCTCCT | Sigma |
| 14[47]12[48]R1X5 | AACAAGAGGGATAAAAATTTTTAGCATAAAGC TT TCCTCCTCCTCCTCCTCCT | Sigma |
| 10[47]8[48]R1X5 | CTGTAGCTTGACTATTATAGTCAGTTCATTGA TT TCCTCCTCCTCCTCCTCCT | Sigma |
| 6[47]4[48]R1X5 | TACGTTAAAGTAATCTTGACAAGAACCGAACT TT TCCTCCTCCTCCTCCTCCT | Sigma |
| 22[79]20[80]R1X5 | TGGAACAACCGCCTGGCCCTGAGGCCCGCT TT TCCTCCTCCTCCTCCTCCT | Sigma |
| 18[79]16[80]R1X5 | GATGTGCTTCAGGAAGATCGCACAATGTGA TT TCCTCCTCCTCCTCCTCCT | Sigma |
| 14[79]12[80]R1X5 | GCTATCAGAAATGCAATGCCTGAATTAGCA TT TCCTCCTCCTCCTCCTCCT | Sigma |
| 10[79]8[80]R1X5 | GATGGCTTATCAAAAAGATTAAGAGCGTCC TT TCCTCCTCCTCCTCCTCCT | Sigma |
| 6[79]4[80]R1X5 | TTATACCACCAAATCAACGTAACGAACGAG TT TCCTCCTCCTCCTCCTCCT | Sigma |
| 22[111]20[112]R1X5 | GCCCGAGAGTCCACGCTGGTTTGCAGCTAACT TT TCCTCCTCCTCCTCCTCCT | Sigma |
| 18[111]16[112]R1X5 | TCTTCGCTGCACCGCTTCTGGTGCGGCCTTCC TT TCCTCCTCCTCCTCCTCCT | Sigma |
| 10[111]8[112]R1X5 | TTGCTCCTTTCAAATATCGCGTTTGAGGGGGT TT TCCTCCTCCTCCTCCTCCT | Sigma |
| 6[111]4[112]R1X5 | ATTACCTTTGAATAAGGCTTGCCCAAATCCGC TT TCCTCCTCCTCCTCCTCCT | Sigma |
| 19[160]20[144]R1X5 | GCAATTCACATATTCCTGATTATCAAAGTGTA TT TCCTCCTCCTCCTCCTCCT | Sigma |
| 15[160]16[144]R1X5 | ATCGCAAGTATGTAAATGCTGATGATAGGAAC TT TCCTCCTCCTCCTCCTCCT | Sigma |
| 7[160]8[144]R1X5 | TTATTACGAAGAACTGGCATGATTGCGAGAGG TT TCCTCCTCCTCCTCCTCCT | Sigma |
| 3[160]4[144]R1X5 | TTGACAGGCCACCACCAGAGCCGCGATTTGTA TT TCCTCCTCCTCCTCCTCCT | Sigma |
| 22[175]20[176]R1X5 | ACCTTGCTTGGTCAGTTGGCAAAGAGCGGA TT TCCTCCTCCTCCTCCTCCT | Sigma |

| | | |
|---|---|---|
| 18[175]16[176]R1X5 | CTGAGCAAAAATTAATTACATTTTGGGTTA TT TCCTCCTCCTCCTCCTCCT | Sigma |
| 10[175]8[176]R1X5 | TTAACGTCTAACATAAAAACAGGTAACGGA TT TCCTCCTCCTCCTCCTCCT | Sigma |
| 6[175]4[176]R1X5 | CAGCAAAAGGAAACGTCACCAATGAGCCGC TT TCCTCCTCCTCCTCCTCCT | Sigma |
| 22[207]20[208]R1X5 | AGCCAGCAATTGAGGAAGGTTATCATCATTTT TT TCCTCCTCCTCCTCCTCCT | Sigma |
| 18[207]16[208]R1X5 | CGCGCAGATTACCTTTTTTAATGGGAGAGACT TT TCCTCCTCCTCCTCCTCCT | Sigma |
| 10[207]8[208]R1X5 | ATCCCAATGAGAATTAACTGAACAGTTACCAG TT TCCTCCTCCTCCTCCTCCT | Sigma |
| 6[207]4[208]R1X5 | TCACCGACGCACCGTAATCAGTAGCAGAACCG TT TCCTCCTCCTCCTCCTCCT | Sigma |
| 22[239]20[240]R1X5 | TTAACACCAGCACTAACAACTAATCGTTATTA TT TCCTCCTCCTCCTCCTCCT | Sigma |
| 18[239]16[240]R1X5 | CCTGATTGCAATATATGTGAGTGATCAATAGT TT TCCTCCTCCTCCTCCTCCT | Sigma |
| 14[239]12[240]R1X5 | AGTATAAAGTTCAGCTAATGCAGATGTCTTTC TT TCCTCCTCCTCCTCCTCCT | Sigma |
| 10[239]8[240]R1X5 | GCCAGTTAGAGGGTAATTGAGCGCTTTAAGAA TT TCCTCCTCCTCCTCCTCCT | Sigma |
| 6[239]4[240]R1X5 | GAAATTATTGCCTTTAGCGTCAGACCGGAACC TT TCCTCCTCCTCCTCCTCCT | Sigma |
| 22[271]20[272]R1X5 | CAGAAGATTAGATAATACATTTGTCGACAA TT TCCTCCTCCTCCTCCTCCT | Sigma |
| 18[271]16[272]R1X5 | CTTTTACAAAATCGTCGCTATTAGCGATAG TT TCCTCCTCCTCCTCCTCCT | Sigma |
| 14[271]12[272]R1X5 | TTAGTATCACAATAGATAAGTCCACGAGCA TT TCCTCCTCCTCCTCCTCCT | Sigma |
| 10[271]8[272]R1X5 | ACGCTAACACCCACAAGAATTGAAAATAGC TT TCCTCCTCCTCCTCCTCCT | Sigma |
| 6[271]4[272]R1X5 | ACCGATTGTCGGCATTTTCGGTCATAATCA TT TCCTCCTCCTCCTCCTCCT | Sigma |
| 21[32]23[31]BLK | TTTTCACTCAAAGGGCGAAAAACCATCACC | Sigma |
| 0[47]1[31]BLLK | AGAAAGGAACAACTAAAGGAATTCAAAAAAA | Sigma |
| 0[111]1[95]BLK | TAAATGAATTTTCTGTATGGGATTAATTTCTT | Sigma |
| 21[128]23[127]BLK | CCCAGCAGGCGAAAAATCCCTTATAAATCAAGCCGGCG | Sigma |
| 2[271]0[272]BLK | GTTTTAACTTAGTACCGCCACCCAGAGCCA | Sigma |
| **Staples with R5×5 extensions for 'Box'-shaped grid** | | |
| 22[47]20[48]R5X5 | CTCCAACGCAGTGAGACGGGCAACCAGCTGCA TT CTTCTTCTTCTTCTTCTTC | Sigma |
| 18[47]16[48]R5X5 | CCAGGGTTGCCAGTTTGAGGGGACCCGTGGGA TT CTTCTTCTTCTTCTTCTTC | Sigma |
| 14[47]12[48]R5X5 | AACAAGAGGGATAAAAATTTTTAGCATAAAGC TT CTTCTTCTTCTTCTTCTTC | Sigma |
| 10[47]8[48]R5X5 | CTGTAGCTTGACTATTATAGTCAGTTCATTGA TT CTTCTTCTTCTTCTTCTTC | Sigma |
| 6[47]4[48]R5X5 | TACGTTAAAGTAATCTTGACAAGAACCGAACT TT CTTCTTCTTCTTCTTCTTC | Sigma |
| 22[79]20[80]R5X5 | TGGAACAACCGCCTGGCCCTGAGGCCCGCT TT CTTCTTCTTCTTCTTCTTC | Sigma |
| 18[79]16[80]R5X5 | GATGTGCTTCAGGAAGATCGCACAATGTGA TT CTTCTTCTTCTTCTTCTTC | Sigma |
| 14[79]12[80]R5X5 | GCTATCAGAAATGCAATGCCTGAATTAGCA TT CTTCTTCTTCTTCTTCTTC | Sigma |
| 10[79]8[80]R5X5 | GATGGCTTATCAAAAAGATTAAGAGCGTCC TT CTTCTTCTTCTTCTTCTTC | Sigma |
| 6[79]4[80]R5X5 | TTATACCACCAAATCAACGTAACGAACGAG TT CTTCTTCTTCTTCTTCTTC | Sigma |
| 22[111]20[112]R5X5 | GCCCGAGAGTCCACGCTGGTTTGCAGCTAACT TT CTTCTTCTTCTTCTTCTTC | Sigma |
| 18[111]16[112]R5X5 | TCTTCGCTGCACCGCTTCTGGTGCGGCCTTCC TT CTTCTTCTTCTTCTTCTTC | Sigma |
| 10[111]8[112]R5X5 | TTGCTCCTTTCAAATATCGCGTTTGAGGGGGT TT CTTCTTCTTCTTCTTCTTC | Sigma |
| 6[111]4[112]R5X5 | ATTACCTTTGAATAAGGCTTGCCCAAATCCGC TT CTTCTTCTTCTTCTTCTTC | Sigma |
| 19[160]20[144]R5X5 | GCAATTCACATATTCCTGATTATCAAAGTGTA TT CTTCTTCTTCTTCTTCTTC | Sigma |
| 15[160]16[144]R5X5 | ATCGCAAGTATGTAAATGCTGATGATAGGAAC TT CTTCTTCTTCTTCTTCTTC | Sigma |
| 7[160]8[144]R5X5 | TTATTACGAAGAACTGGCATGATTGCGAGAGG TT CTTCTTCTTCTTCTTCTTC | Sigma |
| 3[160]4[144]R5X5 | TTGACAGGCCACCACCAGAGCCGCGATTTGTA TT CTTCTTCTTCTTCTTCTTC | Sigma |
| 22[175]20[176]R5X5 | ACCTTGCTTGGTCAGTTGGCAAAGAGCGGA TT CTTCTTCTTCTTCTTCTTC | Sigma |
| 18[175]16[176]R5X5 | CTGAGCAAAAATTAATTACATTTTGGGTTA TT CTTCTTCTTCTTCTTCTTC | Sigma |
| 10[175]8[176]R5X5 | TTAACGTCTAACATAAAAACAGGTAACGGA TT CTTCTTCTTCTTCTTCTTC | Sigma |

| | | |
|---|---|---|
| 6[175]4[176]R5X5 | CAGCAAAAGGAAACGTCACCAATGAGCCGC TT CTTCTTCTTCTTCTTCTTC | Sigma |
| 22[207]20[208]R5X5 | AGCCAGCAATTGAGGAAGGTTATCATCATTTT TT CTTCTTCTTCTTCTTCTTC | Sigma |
| 18[207]16[208]R5X5 | CGCGCAGATTACCTTTTTTAATGGGAGAGACT TT CTTCTTCTTCTTCTTCTTC | Sigma |
| 10[207]8[208]R5X5 | ATCCCAATGAGAATTAACTGAACAGTTACCAG TT CTTCTTCTTCTTCTTCTTC | Sigma |
| 6[207]4[208]R5X5 | TCACCGACGCACCGTAATCAGTAGCAGAACCG TT CTTCTTCTTCTTCTTCTTC | Sigma |
| 22[239]20[240]R5X5 | TTAACACCAGCACTAACAACTAATCGTTATTA TT CTTCTTCTTCTTCTTCTTC | Sigma |
| 18[239]16[240]R5X5 | CCTGATTGCAATATATGTGAGTGATCAATAGT TT CTTCTTCTTCTTCTTCTTC | Sigma |
| 14[239]12[240]R5X5 | AGTATAAAGTTCAGCTAATGCAGATGTCTTTC TT CTTCTTCTTCTTCTTCTTC | Sigma |
| 10[239]8[240]R5X5 | GCCAGTTAGAGGGTAATTGAGCGCTTTAAGAA TT CTTCTTCTTCTTCTTCTTC | Sigma |
| 6[239]4[240]R5X5 | GAAATTATTGCCTTTAGCGTCAGACCGGAACC TT CTTCTTCTTCTTCTTCTTC | Sigma |
| 22[271]20[272]R5X5 | CAGAAGATTAGATAATACATTTGTCGACAA TT CTTCTTCTTCTTCTTCTTC | Sigma |
| 18[271]16[272]R5X5 | CTTTTACAAAATCGTCGCTATTAGCGATAG TT CTTCTTCTTCTTCTTCTTC | Sigma |
| 14[271]12[272]R5X5 | TTAGTATCACAATAGATAAGTCCACGAGCA TT CTTCTTCTTCTTCTTCTTC | Sigma |
| 10[271]8[272]R5X5 | ACGCTAACACCCACAAGAATTGAAAATAGC TT CTTCTTCTTCTTCTTCTTC | Sigma |
| 6[271]4[272]R5X5 | ACCGATTGTCGGCATTTTCGGTCATAATCA TT CTTCTTCTTCTTCTTCTTC | Sigma |
| **Staples with R1×5 extensions for 'C'-shaped grid** | | |
| 22[47]20[48]R1X5 | CTCCAACGCAGTGAGACGGGCAACCAGCTGCA TT TCCTCCTCCTCCTCCTCCT | Sigma |
| 18[47]16[48]R1X5 | CCAGGGTTGCCAGTTTGAGGGGACCCGTGGGA TT TCCTCCTCCTCCTCCTCCT | Sigma |
| 14[47]12[48]R1X5 | AACAAGAGGGATAAAAATTTTTAGCATAAAGC TT TCCTCCTCCTCCTCCTCCT | Sigma |
| 10[47]8[48]R1X5 | CTGTAGCTTGACTATTATAGTCAGTTCATTGA TT TCCTCCTCCTCCTCCTCCT | Sigma |
| 6[47]4[48]R1X5 | TACGTTAAAGTAATCTTGACAAGAACCGAACT TT TCCTCCTCCTCCTCCTCCT | Sigma |
| 22[79]20[80]R1X5 | TGGAACAACCGCCTGGCCCTGAGGCCCGCT TT TCCTCCTCCTCCTCCTCCT | Sigma |
| 18[79]16[80]R1X5 | GATGTGCTTCAGGAAGATCGCACAATGTGA TT TCCTCCTCCTCCTCCTCCT | Sigma |
| 14[79]12[80]R1X5 | GCTATCAGAAATGCAATGCCTGAATTAGCA TT TCCTCCTCCTCCTCCTCCT | Sigma |
| 10[79]8[80]R1X5 | GATGGCTTATCAAAAAGATTAAGAGCGTCC TT TCCTCCTCCTCCTCCTCCT | Sigma |
| 6[79]4[80]R1X5 | TTATACCACCAAATCAACGTAACGAACGAG TT TCCTCCTCCTCCTCCTCCT | Sigma |
| 22[111]20[112]R1X5 | GCCCGAGAGTCCACGCTGGTTTGCAGCTAACT TT TCCTCCTCCTCCTCCTCCT | Sigma |
| 19[160]20[144]R1X5 | GCAATTCACATATTCCTGATTATCAAAGTGTA TT TCCTCCTCCTCCTCCTCCT | Sigma |
| 22[175]20[176]R1X5 | ACCTTGCTTGGTCAGTTGGCAAAGAGCGGA TT TCCTCCTCCTCCTCCTCCT | Sigma |
| 22[207]20[208]R1X5 | AGCCAGCAATTGAGGAAGGTTATCATCATTTT TT TCCTCCTCCTCCTCCTCCT | Sigma |
| 22[239]20[240]R1X5 | TTAACACCAGCACTAACAACTAATCGTTATTA TT TCCTCCTCCTCCTCCTCCT | Sigma |
| 18[239]16[240]R1X5 | CCTGATTGCAATATATGTGAGTGATCAATAGT TT TCCTCCTCCTCCTCCTCCT | Sigma |
| 14[239]12[240]R1X5 | AGTATAAAGTTCAGCTAATGCAGATGTCTTTC TT TCCTCCTCCTCCTCCTCCT | Sigma |
| 10[239]8[240]R1X5 | GCCAGTTAGAGGGTAATTGAGCGCTTTAAGAA TT TCCTCCTCCTCCTCCTCCT | Sigma |
| 6[239]4[240]R1X5 | GAAATTATTGCCTTTAGCGTCAGACCGGAACC TT TCCTCCTCCTCCTCCTCCT | Sigma |
| 22[271]20[272]R1X5 | CAGAAGATTAGATAATACATTTGTCGACAA TT TCCTCCTCCTCCTCCTCCT | Sigma |
| 18[271]16[272]R1X5 | CTTTTACAAAATCGTCGCTATTAGCGATAG TT TCCTCCTCCTCCTCCTCCT | Sigma |
| 14[271]12[272]R1X5 | TTAGTATCACAATAGATAAGTCCACGAGCA TT TCCTCCTCCTCCTCCTCCT | Sigma |
| 10[271]8[272]R1X5 | ACGCTAACACCCACAAGAATTGAAAATAGC TT TCCTCCTCCTCCTCCTCCT | Sigma |
| 6[271]4[272]R1X5 | ACCGATTGTCGGCATTTTCGGTCATAATCA TT TCCTCCTCCTCCTCCTCCT | Sigma |
| **Staples with R5×5 extensions for 'C'-shaped grid** | | |
| 22[47]20[48]R5X5 | CTCCAACGCAGTGAGACGGGCAACCAGCTGCA TT CTTCTTCTTCTTCTTCTTC | Sigma |
| 18[47]16[48]R5X5 | CCAGGGTTGCCAGTTTGAGGGGACCCGTGGGA TT CTTCTTCTTCTTCTTCTTC | Sigma |
| 14[47]12[48]R5X5 | AACAAGAGGGATAAAAATTTTTAGCATAAAGC TT CTTCTTCTTCTTCTTCTTC | Sigma |

| | | |
|---|---|---|
| 10[47]8[48]R5X5 | CTGTAGCTTGACTATTATAGTCAGTTCATTGA TT CTTCTTCTTCTTCTTCTTC | Sigma |
| 6[47]4[48]R5X5 | TACGTTAAAGTAATCTTGACAAGAACCGAACT TT CTTCTTCTTCTTCTTCTTC | Sigma |
| 22[79]20[80]R5X5 | TGGAACAACCGCCTGGCCCTGAGGCCCGCT TT CTTCTTCTTCTTCTTCTTC | Sigma |
| 18[79]16[80]R5X5 | GATGTGCTTCAGGAAGATCGCACAATGTGA TT CTTCTTCTTCTTCTTCTTC | Sigma |
| 14[79]12[80]R5X5 | GCTATCAGAAATGCAATGCCTGAATTAGCA TT CTTCTTCTTCTTCTTCTTC | Sigma |
| 10[79]8[80]R5X5 | GATGGCTTATCAAAAAGATTAAGAGCGTCC TT CTTCTTCTTCTTCTTCTTC | Sigma |
| 6[79]4[80]R5X5 | TTATACCACCAAATCAACGTAACGAACGAG TT CTTCTTCTTCTTCTTCTTC | Sigma |
| 22[111]20[112]R5X5 | GCCCGAGAGTCCACGCTGGTTTGCAGCTAACT TT CTTCTTCTTCTTCTTCTTC | Sigma |
| 19[160]20[144]R5X5 | GCAATTCACATATTCCTGATTATCAAAGTGTA TT CTTCTTCTTCTTCTTCTTC | Sigma |
| 22[175]20[176]R5X5 | ACCTTGCTTGGTCAGTTGGCAAAGAGCGGA TT CTTCTTCTTCTTCTTCTTC | Sigma |
| 22[207]20[208]R5X5 | AGCCAGCAATTGAGGAAGGTTATCATCATTTT TT CTTCTTCTTCTTCTTCTTC | Sigma |
| 22[239]20[240]R5X5 | TTAACACCAGCACTAACAACTAATCGTTATTA TT CTTCTTCTTCTTCTTCTTC | Sigma |
| 18[239]16[240]R5X5 | CCTGATTGCAATATATGTGAGTGATCAATAGT TT CTTCTTCTTCTTCTTCTTC | Sigma |
| 14[239]12[240]R5X5 | AGTATAAAGTTCAGCTAATGCAGATGTCTTTC TT CTTCTTCTTCTTCTTCTTC | Sigma |
| 10[239]8[240]R5X5 | GCCAGTTAGAGGGTAATTGAGCGCTTTAAGAA TT CTTCTTCTTCTTCTTCTTC | Sigma |
| 6[239]4[240]R5X5 | GAAATTATTGCCTTTAGCGTCAGACCGGAACC TT CTTCTTCTTCTTCTTCTTC | Sigma |
| 22[271]20[272]R5X5 | CAGAAGATTAGATAATACATTTGTCGACAA TT CTTCTTCTTCTTCTTCTTC | Sigma |
| 18[271]16[272]R5X5 | CTTTTACAAAATCGTCGCTATTAGCGATAG TT CTTCTTCTTCTTCTTCTTC (Not used in Barcoding) | Sigma |
| 14[271]12[272]R5X5 | TTAGTATCACAATAGATAAGTCCACGAGCA TT CTTCTTCTTCTTCTTCTTC | Sigma |
| 10[271]8[272]R5X5 | ACGCTAACACCCACAAGAATTGAAAATAGC TT CTTCTTCTTCTTCTTCTTC | Sigma |
| 6[271]4[272]R5X5 | ACCGATTGTCGGCATTTTCGGTCATAATCA TT CTTCTTCTTCTTCTTCTTC | Sigma |
| **Staples with R1×5 extensions for 'U'-shaped grid** | | |
| 22[47]20[48]R1X5 | CTCCAACGCAGTGAGACGGGCAACCAGCTGCA TT TCCTCCTCCTCCTCCTCCT | Sigma |
| 18[47]16[48]R1X5 | CCAGGGTTGCCAGTTTGAGGGGACCCGTGGGA TT TCCTCCTCCTCCTCCTCCT | Sigma |
| 14[47]12[48]R1X5 | AACAAGAGGGATAAAAATTTTTAGCATAAAGC TT TCCTCCTCCTCCTCCTCCT | Sigma |
| 10[47]8[48]R1X5 | CTGTAGCTTGACTATTATAGTCAGTTCATTGA TT TCCTCCTCCTCCTCCTCCT | Sigma |
| 6[47]4[48]R1X5 | TACGTTAAAGTAATCTTGACAAGAACCGAACT TT TCCTCCTCCTCCTCCTCCT | Sigma |
| 22[79]20[80]R1X5 | TGGAACAACCGCCTGGCCCTGAGGCCCGCT TT TCCTCCTCCTCCTCCTCCT | Sigma |
| 18[79]16[80]R1X5 | GATGTGCTTCAGGAAGATCGCACAATGTGA TT TCCTCCTCCTCCTCCTCCT | Sigma |
| 14[79]12[80]R1X5 | GCTATCAGAAATGCAATGCCTGAATTAGCA TT TCCTCCTCCTCCTCCTCCT | Sigma |
| 10[79]8[80]R1X5 | GATGGCTTATCAAAAAGATTAAGAGCGTCC TT TCCTCCTCCTCCTCCTCCT | Sigma |
| 6[79]4[80]R1X5 | TTATACCACCAAATCAACGTAACGAACGAG TT TCCTCCTCCTCCTCCTCCT | Sigma |
| 22[111]20[112]R1X5 | GCCCGAGAGTCCACGCTGGTTTGCAGCTAACT TT TCCTCCTCCTCCTCCTCCT | Sigma |
| 6[111]4[112]R1X5 | ATTACCTTTGAATAAGGCTTGCCCAAATCCGC TT TCCTCCTCCTCCTCCTCCT | Sigma |
| 19[160]20[144]R1X5 | GCAATTCACATATTCCTGATTATCAAAGTGTA TT TCCTCCTCCTCCTCCTCCT | Sigma |
| 3[160]4[144]R1X5 | TTGACAGGCCACCACCAGAGCCGCGATTTGTA TT TCCTCCTCCTCCTCCTCCT | Sigma |
| 22[175]20[176]R1X5 | ACCTTGCTTGGTCAGTTGGCAAAGAGCGGA TT TCCTCCTCCTCCTCCTCCT | Sigma |
| 6[175]4[176]R1X5 | CAGCAAAAGGAAACGTCACCAATGAGCCGC TT TCCTCCTCCTCCTCCTCCT | Sigma |
| 22[207]20[208]R1X5 | AGCCAGCAATTGAGGAAGGTTATCATCATTTT TT TCCTCCTCCTCCTCCTCCT | Sigma |
| 6[207]4[208]R1X5 | TCACCGACGCACCGTAATCAGTAGCAGAACCG TT TCCTCCTCCTCCTCCTCCT | Sigma |
| 22[239]20[240]R1X5 | TTAACACCAGCACTAACAACTAATCGTTATTA TT TCCTCCTCCTCCTCCTCCT | Sigma |
| 6[239]4[240]R1X5 | GAAATTATTGCCTTTAGCGTCAGACCGGAACC TT TCCTCCTCCTCCTCCTCCT | Sigma |
| 22[271]20[272]R1X5 | CAGAAGATTAGATAATACATTTGTCGACAA TT TCCTCCTCCTCCTCCTCCT | Sigma |
| 6[271]4[272]R1X5 | ACCGATTGTCGGCATTTTCGGTCATAATCA TT TCCTCCTCCTCCTCCTCCT | Sigma |

| | | |
|---|---|---|
| **Staples with R5×5 extensions for 'U'-shaped grid** | | |
| 22[47]20[48]R5X5 | CTCCAACGCAGTGAGACGGGCAACCAGCTGCA TT CTTCTTCTTCTTCTTCTTC | Sigma |
| 18[47]16[48]R5X5 | CCAGGGTTGCCAGTTTGAGGGGACCCGTGGGA TT CTTCTTCTTCTTCTTCTTC | Sigma |
| 14[47]12[48]R5X5 | AACAAGAGGGATAAAAATTTTTAGCATAAAGC TT CTTCTTCTTCTTCTTCTTC | Sigma |
| 10[47]8[48]R5X5 | CTGTAGCTTGACTATTATAGTCAGTTCATTGA TT CTTCTTCTTCTTCTTCTTC | Sigma |
| 6[47]4[48]R5X5 | TACGTTAAAGTAATCTTGACAAGAACCGAACT TT CTTCTTCTTCTTCTTCTTC | Sigma |
| 22[79]20[80]R5X5 | TGGAACAACCGCCTGGCCCTGAGGCCCGCT TT CTTCTTCTTCTTCTTCTTC | Sigma |
| 18[79]16[80]R5X5 | GATGTGCTTCAGGAAGATCGCACAATGTGA TT CTTCTTCTTCTTCTTCTTC | Sigma |
| 14[79]12[80]R5X5 | GCTATCAGAAATGCAATGCCTGAATTAGCA TT CTTCTTCTTCTTCTTCTTC | Sigma |
| 10[79]8[80]R5X5 | GATGGCTTATCAAAAAGATTAAGAGCGTCC TT CTTCTTCTTCTTCTTCTTC | Sigma |
| 6[79]4[80]R5X5 | TTATACCACCAAATCAACGTAACGAACGAG TT CTTCTTCTTCTTCTTCTTC | Sigma |
| 22[111]20[112]R5X5 | GCCCGAGAGTCCACGCTGGTTTGCAGCTAACT TT CTTCTTCTTCTTCTTCTTC | Sigma |
| 6[111]4[112]R5X5 | ATTACCTTTGAATAAGGCTTGCCCAAATCCGC TT CTTCTTCTTCTTCTTCTTC | Sigma |
| 19[160]20[144]R5X5 | GCAATTCACATATTCCTGATTATCAAAGTGTA TT CTTCTTCTTCTTCTTCTTC | Sigma |
| 3[160]4[144]R5X5 | TTGACAGGCCACCACCAGAGCCGCGATTTGTA TT CTTCTTCTTCTTCTTCTTC | Sigma |
| 22[175]20[176]R5X5 | ACCTTGCTTGGTCAGTTGGCAAAGAGCGGA TT CTTCTTCTTCTTCTTCTTC | Sigma |
| 6[175]4[176]R5X5 | CAGCAAAAGGAAACGTCACCAATGAGCCGC TT CTTCTTCTTCTTCTTCTTC | Sigma |
| 22[207]20[208]R5X5 | AGCCAGCAATTGAGGAAGGTTATCATCATTTT TT CTTCTTCTTCTTCTTCTTC | Sigma |
| 6[207]4[208]R5X5 | TCACCGACGCACCGTAATCAGTAGCAGAACCG TT CTTCTTCTTCTTCTTCTTC | Sigma |
| 22[239]20[240]R5X5 | TTAACACCAGCACTAACAACTAATCGTTATTA TT CTTCTTCTTCTTCTTCTTC | Sigma |
| 6[239]4[240]R5X5 | GAAATTATTGCCTTTAGCGTCAGACCGGAACC TT CTTCTTCTTCTTCTTCTTC | Sigma |
| 22[271]20[272]R5X5 | CAGAAGATTAGATAATACATTTGTCGACAA TT CTTCTTCTTCTTCTTCTTC | Sigma |
| 6[271]4[272]R5X5 | ACCGATTGTCGGCATTTTCGGTCATAATCA TT CTTCTTCTTCTTCTTCTTC | Sigma |
| **Staples with R1×5 extensions for 'H'-shaped grid** | | |
| 22[47]20[48]R1X5 | CTCCAACGCAGTGAGACGGGCAACCAGCTGCA TT TCCTCCTCCTCCTCCTCCT | Sigma |
| 6[47]4[48]R1X5 | TACGTTAAAGTAATCTTGACAAGAACCGAACT TT TCCTCCTCCTCCTCCTCCT | Sigma |
| 22[79]20[80]R1X5 | TGGAACAACCGCCTGGCCCTGAGGCCCGCT TT TCCTCCTCCTCCTCCTCCT | Sigma |
| 6[79]4[80]R1X5 | TTATACCACCAAATCAACGTAACGAACGAG TT TCCTCCTCCTCCTCCTCCT | Sigma |
| 22[111]20[112]R1X5 | GCCCGAGAGTCCACGCTGGTTTGCAGCTAACT TT TCCTCCTCCTCCTCCTCCT | Sigma |
| 6[111]4[112]R1X5 | ATTACCTTTGAATAAGGCTTGCCCAAATCCGC TT TCCTCCTCCTCCTCCTCCT | Sigma |
| 19[160]20[144]R1X5 | GCAATTCACATATTCCTGATTATCAAAGTGTA TT TCCTCCTCCTCCTCCTCCT | Sigma |
| 15[160]16[144]R1X5 | ATCGCAAGTATGTAAATGCTGATGATAGGAAC TT TCCTCCTCCTCCTCCTCCT | Sigma |
| 7[160]8[144]R1X5 | TTATTACGAAGAACTGGCATGATTGCGAGAGG TT TCCTCCTCCTCCTCCTCCT | Sigma |
| 3[160]4[144]R1X5 | TTGACAGGCCACCACCAGAGCCGCGATTTGTA TT TCCTCCTCCTCCTCCTCCT | Sigma |
| 22[175]20[176]R1X5 | ACCTTGCTTGGTCAGTTGGCAAAGAGCGGA TT TCCTCCTCCTCCTCCTCCT | Sigma |
| 18[175]16[176]R1X5 | CTGAGCAAAAATTAATTACATTTTGGGTTA TT TCCTCCTCCTCCTCCTCCT | Sigma |
| 10[175]8[176]R1X5 | TTAACGTCTAACATAAAAACAGGTAACGGA TT TCCTCCTCCTCCTCCTCCT | Sigma |
| 6[175]4[176]R1X5 | CAGCAAAAGGAAACGTCACCAATGAGCCGC TT TCCTCCTCCTCCTCCTCCT | Sigma |
| 22[207]20[208]R1X5 | AGCCAGCAATTGAGGAAGGTTATCATCATTTT TT TCCTCCTCCTCCTCCTCCT | Sigma |
| 6[207]4[208]R1X5 | TCACCGACGCACCGTAATCAGTAGCAGAACCG TT TCCTCCTCCTCCTCCTCCT | Sigma |
| 22[239]20[240]R1X5 | TTAACACCAGCACTAACAACTAATCGTTATTA TT TCCTCCTCCTCCTCCTCCT | Sigma |
| 6[239]4[240]R1X5 | GAAATTATTGCCTTTAGCGTCAGACCGGAACC TT TCCTCCTCCTCCTCCTCCT | Sigma |
| 22[271]20[272]R1X5 | CAGAAGATTAGATAATACATTTGTCGACAA TT TCCTCCTCCTCCTCCTCCT | Sigma |
| 6[271]4[272]R1X5 | ACCGATTGTCGGCATTTTCGGTCATAATCA TT TCCTCCTCCTCCTCCTCCT | Sigma |

| Staples with R5×5 extensions for 'H'-shaped grid | | |
|---|---|---|
| 22[47]20[48]R5X5 | CTCCAACGCAGTGAGACGGGCAACCAGCTGCA TT CTTCTTCTTCTTCTTCTTC | Sigma |
| 6[47]4[48]R5X5 | TACGTTAAAGTAATCTTGACAAGAACCGAACT TT CTTCTTCTTCTTCTTCTTC | Sigma |
| 22[79]20[80]R5X5 | TGGAACAACCGCCTGGCCCTGAGGCCCGCT TT CTTCTTCTTCTTCTTCTTC | Sigma |
| 6[79]4[80]R5X5 | TTATACCACCAAATCAACGTAACGAACGAG TT CTTCTTCTTCTTCTTCTTC | Sigma |
| 22[111]20[112]R5X5 | GCCCGAGAGTCCACGCTGGTTTGCAGCTAACT TT CTTCTTCTTCTTCTTCTTC | Sigma |
| 6[111]4[112]R5X5 | ATTACCTTTGAATAAGGCTTGCCCAAATCCGC TT CTTCTTCTTCTTCTTCTTC | Sigma |
| 19[160]20[144]R5X5 | GCAATTCACATATTCCTGATTATCAAAGTGTA TT CTTCTTCTTCTTCTTCTTC | Sigma |
| 15[160]16[144]R5X5 | ATCGCAAGTATGTAAATGCTGATGATAGGAAC TT CTTCTTCTTCTTCTTCTTC | Sigma |
| 7[160]8[144]R5X5 | TTATTACGAAGAACTGGCATGATTGCGAGAGG TT CTTCTTCTTCTTCTTCTTC | Sigma |
| 3[160]4[144]R5X5 | TTGACAGGCCACCACCAGAGCCGCGATTTGTA TT CTTCTTCTTCTTCTTCTTC | Sigma |
| 22[175]20[176]R5X5 | ACCTTGCTTGGTCAGTTGGCAAAGAGCGGA TT CTTCTTCTTCTTCTTCTTC | Sigma |
| 18[175]16[176]R5X5 | CTGAGCAAAAATTAATTACATTTGGGTTA TT CTTCTTCTTCTTCTTCTTC | Sigma |
| 10[175]8[176]R5X5 | TTAACGTCTAACATAAAAACAGGTAACGGA TT CTTCTTCTTCTTCTTCTTC | Sigma |
| 6[175]4[176]R5X5 | CAGCAAAAGGAAACGTCACCAATGAGCCGC TT CTTCTTCTTCTTCTTCTTC | Sigma |
| 22[207]20[208]R5X5 | AGCCAGCAATTGAGGAAGGTTATCATCATTTT TT CTTCTTCTTCTTCTTCTTC | Sigma |
| 6[207]4[208]R5X5 | TCACCGACGCACCGTAATCAGTAGCAGAACCG TT CTTCTTCTTCTTCTTCTTC | Sigma |
| 22[239]20[240]R5X5 | TTAACACCAGCACTAACAACTAATCGTTATTA TT CTTCTTCTTCTTCTTCTTC | Sigma |
| 6[239]4[240]R5X5 | GAAATTATTGCCTTTAGCGTCAGACCGGAACC TT CTTCTTCTTCTTCTTCTTC | Sigma |
| 22[271]20[272]R5X5 | CAGAAGATTAGATAATACATTTGTCGACAA TT CTTCTTCTTCTTCTTCTTC | Sigma |
| 6[271]4[272]R5X5 | ACCGATTGTCGGCATTTTCGGTCATAATCA TT CTTCTTCTTCTTCTTCTTC | Sigma |
| **Staples with R1×5 extensions for 'L'-shaped grid** | | |
| 22[47]20[48]R1X5 | CTCCAACGCAGTGAGACGGGCAACCAGCTGCA TT TCCTCCTCCTCCTCCTCCT | Sigma |
| 18[47]16[48]R1X5 | CCAGGGTTGCCAGTTTGAGGGGACCCGTGGGA TT TCCTCCTCCTCCTCCTCCT | Sigma |
| 14[47]12[48]R1X5 | AACAAGAGGGATAAAAATTTTTAGCATAAAGC TT TCCTCCTCCTCCTCCTCCT | Sigma |
| 10[47]8[48]R1X5 | CTGTAGCTTGACTATTATAGTCAGTTCATTGA TT TCCTCCTCCTCCTCCTCCT | Sigma |
| 6[47]4[48]R1X5 | TACGTTAAAGTAATCTTGACAAGAACCGAACT TT TCCTCCTCCTCCTCCTCCT | Sigma |
| 22[79]20[80]R1X5 | TGGAACAACCGCCTGGCCCTGAGGCCCGCT TT TCCTCCTCCTCCTCCTCCT | Sigma |
| 18[79]16[80]R1X5 | GATGTGCTTCAGGAAGATCGCACAATGTGA TT TCCTCCTCCTCCTCCTCCT | Sigma |
| 14[79]12[80]R1X5 | GCTATCAGAAATGCAATGCCTGAATTAGCA TT TCCTCCTCCTCCTCCTCCT | Sigma |
| 10[79]8[80]R1X5 | GATGGCTTATCAAAAAGATTAAGAGCGTCC TT TCCTCCTCCTCCTCCTCCT | Sigma |
| 6[79]4[80]R1X5 | TTATACCACCAAATCAACGTAACGAACGAG TT TCCTCCTCCTCCTCCTCCT | Sigma |
| 22[111]20[112]R1X5 | GCCCGAGAGTCCACGCTGGTTTGCAGCTAACT TT TCCTCCTCCTCCTCCTCCT | Sigma |
| 18[111]16[112]R1X5 | TCTTCGCTGCACCGCTTCTGGTGCGGCCTTCC TT TCCTCCTCCTCCTCCTCCT | Sigma |
| 19[160]20[144]R1X5 | GCAATTCACATATTCCTGATTATCAAAGTGTA TT TCCTCCTCCTCCTCCTCCT | Sigma |
| 15[160]16[144]R1X5 | ATCGCAAGTATGTAAATGCTGATGATAGGAAC TT TCCTCCTCCTCCTCCTCCT | Sigma |
| 22[175]20[176]R1X5 | ACCTTGCTTGGTCAGTTGGCAAAGAGCGGA TT TCCTCCTCCTCCTCCTCCT | Sigma |
| 18[175]16[176]R1X5 | CTGAGCAAAAATTAATTACATTTGGGTTA TT TCCTCCTCCTCCTCCTCCT | Sigma |
| 22[207]20[208]R1X5 | AGCCAGCAATTGAGGAAGGTTATCATCATTTT TT TCCTCCTCCTCCTCCTCCT | Sigma |
| 18[207]16[208]R1X5 | CGCGCAGATTACCTTTTTTAATGGGAGAGACT TT TCCTCCTCCTCCTCCTCCT | Sigma |
| 22[239]20[240]R1X5 | TTAACACCAGCACTAACAACTAATCGTTATTA TT TCCTCCTCCTCCTCCTCCT | Sigma |
| 18[239]16[240]R1X5 | CCTGATTGCAATATATGTGAGTGATCAATAGT TT TCCTCCTCCTCCTCCTCCT | Sigma |
| 22[271]20[272]R1X5 | CAGAAGATTAGATAATACATTTGTCGACAA TT TCCTCCTCCTCCTCCTCCT | Sigma |
| 18[271]16[272]R1X5 | CTTTTACAAAATCGTCGCTATTAGCGATAG TT TCCTCCTCCTCCTCCTCCT | Sigma |

| Staples with R5×5 extensions for 'L'-shaped grid | | |
|---|---|---|
| 22[47]20[48]R5X5 | CTCCAACGCAGTGAGACGGGCAACCAGCTGCA TT CTTCTTCTTCTTCTTCTTC | Sigma |
| 18[47]16[48]R5X5 | CCAGGGTTGCCAGTTTGAGGGGACCCGTGGGA TT CTTCTTCTTCTTCTTCTTC | Sigma |
| 14[47]12[48]R5X5 | AACAAGAGGGATAAAAATTTTTAGCATAAAGC TT CTTCTTCTTCTTCTTCTTC | Sigma |
| 10[47]8[48]R5X5 | CTGTAGCTTGACTATTATAGTCAGTTCATTGA TT CTTCTTCTTCTTCTTCTTC | Sigma |
| 6[47]4[48]R5X5 | TACGTTAAAGTAATCTTGACAAGAACCGAACT TT CTTCTTCTTCTTCTTCTTC | Sigma |
| 22[79]20[80]R5X5 | TGGAACAACCGCCTGGCCCTGAGGCCCGCT TT CTTCTTCTTCTTCTTCTTC | Sigma |
| 18[79]16[80]R5X5 | GATGTGCTTCAGGAAGATCGCACAATGTGA TT CTTCTTCTTCTTCTTCTTC | Sigma |
| 14[79]12[80]R5X5 | GCTATCAGAAATGCAATGCCTGAATTAGCA TT CTTCTTCTTCTTCTTCTTC | Sigma |
| 10[79]8[80]R5X5 | GATGGCTTATCAAAAAGATTAAGAGCGTCC TT CTTCTTCTTCTTCTTCTTC | Sigma |
| 6[79]4[80]R5X5 | TTATACCACCAAATCAACGTAACGAACGAG TT CTTCTTCTTCTTCTTCTTC | Sigma |
| 22[111]20[112]R5X5 | GCCCGAGAGTCCACGCTGGTTTGCAGCTAACT TT CTTCTTCTTCTTCTTCTTC | Sigma |
| 18[111]16[112]R5X5 | TCTTCGCTGCACCGCTTCTGGTGCGGCCTTCC TT CTTCTTCTTCTTCTTCTTC | Sigma |
| 19[160]20[144]R5X5 | GCAATTCACATATTCCTGATTATCAAAGTGTA TT CTTCTTCTTCTTCTTCTTC | Sigma |
| 15[160]16[144]R5X5 | ATCGCAAGTATGTAAATGCTGATGATAGGAAC TT CTTCTTCTTCTTCTTCTTC | Sigma |
| 22[175]20[176]R5X5 | ACCTTGCTTGGTCAGTTGGCAAAGAGCGGA TT CTTCTTCTTCTTCTTCTTC | Sigma |
| 18[175]16[176]R5X5 | CTGAGCAAAAATTAATTACATTTTGGGTTA TT CTTCTTCTTCTTCTTCTTC | Sigma |
| 22[207]20[208]R5X5 | AGCCAGCAATTGAGGAAGGTTATCATCATTTT TT CTTCTTCTTCTTCTTCTTC | Sigma |
| 18[207]16[208]R5X5 | CGCGCAGATTACCTTTTTTAATGGGAGAGACT TT CTTCTTCTTCTTCTTCTTC | Sigma |
| 22[239]20[240]R5X5 | TTAACACCAGCACTAACAACTAATCGTTATTA TT CTTCTTCTTCTTCTTCTTC | Sigma |
| 18[239]16[240]R5X5 | CCTGATTGCAATATATGTGAGTGATCAATAGT TT CTTCTTCTTCTTCTTCTTC | Sigma |
| 22[271]20[272]R5X5 | CAGAAGATTAGATAATACATTTGTCGACAA TT CTTCTTCTTCTTCTTCTTC | Sigma |
| 18[271]16[272]R5X5 | CTTTTACAAAATCGTCGCTATTAGCGATAG TT CTTCTTCTTCTTCTTCTTC | Sigma |

**Supplementary table 6:** Staples with Biotin

| Staple Name | Staple Sequence | Manufacturer |
|---|---|---|
| 18[63]20[56]BIOTIN | [BTN]ATTAAGTTTACCGAGCTCGAATTCGGGAAACCTGTCGTGC | Sigma |
| 4[63]6[56]BIOTIN | [BTN]ATAAGGGAACCGGATATTCATTACGTCAGGACGTTGGGAA | Sigma |
| 18[127]20[120]BIOTIN | [BTN]GCGATCGGCAATTCCACACAACAGGTGCCTAATGAGTG | Sigma |
| 4[127]6[120]BIOTIN | [BTN]TTGTGTCGTGACGAGAAACACCAAATTTCAACTTTAAT | Sigma |
| 18[191]20[184]BIOTIN | [BTN]ATTCATTTTTGTTTGGATTATACTAAGAAACCACCAGAAG | Sigma |
| 4[191]6[184]BIOTIN | [BTN]CACCCTCAGAAACCATCGATAGCATTGAGCCATTTGGGAA | Sigma |
| 18[255]20[248]BIOTIN | [BTN]AACAATAACGTAAAACAGAAATAAAAATCCTTTGCCCGAA | Sigma |
| 4[255]6[248]BIOTIN | [BTN]AGCCACCACTGTAGCGCGTTTTCAAGGGAGGGAAGGTAAA | Sigma |

**Supplementary table 7:** Staples for Stem Localizer and Stacking read out

| Staple Name | Staple Sequence | Manufacturer |
|---|---|---|
| Stem Localizer Staples with R1×5 Docking Strands | | |
| R1x5_12[143]11[159]_TAi | TCCTCCTCCTCCTCCTCCTTTTTCTACTACGCGAGCTGAAAAGGTTACCGCGCAGAAA | Sigma |
| R1x5_12[143]11[159]_ATi | TCCTCCTCCTCCTCCTCCTTTTTCTACTACGCGAGCTGAAAAGGTTACCGCGCAGAAT | Sigma |
| R1x5_12[143]11[159]_CGi | TCCTCCTCCTCCTCCTCCTTTTTCTACTACGCGAGCTGAAAAGGTTACCGCGCAGAAG | Sigma |
| R1x5_12[143]11[159]_GCi | TCCTCCTCCTCCTCCTCCTTTTTCTACTACGCGAGCTGAAAAGGTTACCGCGCAGAAC | Sigma |

| | | |
|---|---|---|
| R1x5_12[143]11[159]_GAP | TCCTCCTCCTCCTCCTCCTTTTTCTACTACGCGAGCTGAAAAGGTTACCGCGCAGA | Sigma |
| **Stem Localizer Staples with R5×5 Docking Strands** | | |
| R5x5_12[143]11[159]_TAi | CTTCTTCTTCTTCTTCTTCTTTTCTACTACGCGAGCTGAAAAGGTTACCGCGCAGAAA | Sigma |
| R5x5_12[143]11[159]_ATi | CTTCTTCTTCTTCTTCTTCTTTTCTACTACGCGAGCTGAAAAGGTTACCGCGCAGAAT | Sigma |
| R5x5_12[143]11[159]_CGi | CTTCTTCTTCTTCTTCTTCTTTTCTACTACGCGAGCTGAAAAGGTTACCGCGCAGAAG | Sigma |
| R5x5_12[143]11[159]_GCi | CTTCTTCTTCTTCTTCTTCTTTTCTACTACGCGAGCTGAAAAGGTTACCGCGCAGAAC | Sigma |
| R5x5_12[143]11[159]_GAP | CTTCTTCTTCTTCTTCTTCTTTTCTACTACGCGAGCTGAAAAGGTTACCGCGCAGA | Sigma |
| **Staples for N\|A Stacks** | | |
| 11[160]12[144]_R1_7_TAi | TCCTCCTTTTCTCCAATAGCTCATCGTAGGAATCATGGCATCAA | Sigma |
| 11[160]12[144]_R1_7_ATi | TCCTCCTATTCTCCAATAGCTCATCGTAGGAATCATGGCATCAA | Sigma |
| 11[160]12[144]_R1_7_CGi | TCCTCCTCTTCTCCAATAGCTCATCGTAGGAATCATGGCATCAA | Sigma |
| 11[160]12[144]_R1_7_GCi | TCCTCCTGTTCTCCAATAGCTCATCGTAGGAATCATGGCATCAA | Sigma |
| **Staples for N\|G Stacks** | | |
| 11[160]12[144]_R5_7_TAi | CTTCTTCTTTCTCCAATAGCTCATCGTAGGAATCATGGCATCAA | Sigma |
| 11[160]12[144]_R5_7_ATi | CTTCTTCATTCTCCAATAGCTCATCGTAGGAATCATGGCATCAA | Sigma |
| 11[160]12[144]_R5_7_CGi | CTTCTTCCTTCTCCAATAGCTCATCGTAGGAATCATGGCATCAA | Sigma |
| 11[160]12[144]_R5_7_GCi | CTTCTTCGTTCTCCAATAGCTCATCGTAGGAATCATGGCATCAA | Sigma |
| **Staples for N\|T Stacks** | | |
| 11[160]12[144]_R4_7_TAi | ACACACATTTCTCCAATAGCTCATCGTAGGAATCATGGCATCAA | Sigma |
| 11[160]12[144]_R4_7_ATi | ACACACAATTCTCCAATAGCTCATCGTAGGAATCATGGCATCAA | Sigma |
| 11[160]12[144]_R4_7_CGi | ACACACACTTCTCCAATAGCTCATCGTAGGAATCATGGCATCAA | Sigma |
| 11[160]12[144]_R4_7_GCi | ACACACAGTTCTCCAATAGCTCATCGTAGGAATCATGGCATCAA | Sigma |
| **Staples for N\|C Stacks** | | |
| 11[160]12[144]_R5_7R_TAi | GAAGAAGTTTCTCCAATAGCTCATCGTAGGAATCATGGCATCAA | Sigma |
| 11[160]12[144]_R5_7R_ATi | GAAGAAGATTCTCCAATAGCTCATCGTAGGAATCATGGCATCAA | Sigma |
| 11[160]12[144]_R5_7R_CGi | GAAGAAGCTTCTCCAATAGCTCATCGTAGGAATCATGGCATCAA | Sigma |
| 11[160]12[144]_R5_7R_GCi | GAAGAAGGTTCTCCAATAGCTCATCGTAGGAATCATGGCATCAA | Sigma |

**Supplementary table 8:** Staples with amine modifications for fluorophore conjugation

| Sequence Name | Sequence | Manufacturer |
|---|---|---|
| R1_7nt | AGGAGGA[AmC3] | Sigma |
| R4_7nt | TGTGTGT[AmC3] | Sigma |
| R5_7nt | GAAGAAG[AmC3] | Sigma |
| R5_7nt_Reverse | CTTCTTC[AmC3] | Sigma |
| Amine_S1_R2x3 | [AmC3]TTTTCCTCTACCACCTACATCACTTACCACCACCACCA | Sigma |

**Supplementary table 9:** Blank Staples specific for Photobleaching and Stem Loop Assay

| Staple Name | Staple Sequence | Manufacturer |
|---|---|---|
| 18[47]16[48]BLK | CCAGGGTTGCCAGTTTGAGGGGACCCGTGGGA BLK | Sigma |
| 10[47]8[48]BLK | CTGTAGCTTGACTATTATAGTCAGTTCATTGA BLK | Sigma |
| 22[79]20[80]BLK | TGGAACAACCGCCTGGCCCTGAGGCCCGCT BLK | Sigma |
| 18[79]16[80]BLK | GATGTGCTTCAGGAAGATCGCACAATGTGA BLK | Sigma |
| 14[79]12[80]BLK | GCTATCAGAAATGCAATGCCTGAATTAGCA BLK | Sigma |

| | | |
|---|---|---|
| 10[79]8[80]BLK | GATGGCTTATCAAAAAGATTAAGAGCGTCC BLK | Sigma |
| 6[79]4[80]BLK | TTATACCACCAAATCAACGTAACGAACGAG BLK | Sigma |
| 18[111]16[112]BLK | TCTTCGCTGCACCGCTTCTGGTGCGGCCTTCC BLK | Sigma |
| 10[111]8[112]BLK | TTGCTCCTTTCAAATATCGCGTTTGAGGGGGT BLK | Sigma |
| 19[160]20[144]BLK | GCAATTCACATATTCCTGATTATCAAAGTGTA BLK | Sigma |
| 15[160]16[144]BLK | ATCGCAAGTATGTAAATGCTGATGATAGGAAC BLK | Sigma |
| 11[160]12[144]BLK | CCAATAGCTCATCGTAGGAATCATGGCATCAA BLK | Sigma |
| 7[160]8[144]BLK | TTATTACGAAGAACTGGCATGATTGCGAGAGG BLK | Sigma |
| 3[160]4[144]BLK | TTGACAGGCCACCACCAGAGCCGCGATTTGTA BLK | Sigma |
| 18[175]16[176]BLK | CTGAGCAAAAATTAATTACATTTTGGGTTA BLK | Sigma |
| 10[175]8[176]BLK | TTAACGTCTAACATAAAAACAGGTAACGGA BLK | Sigma |
| 22[207]20[208]BLK | AGCCAGCAATTGAGGAAGGTTATCATCATTTT BLK | Sigma |
| 18[207]16[208]BLK | CGCGCAGATTACCTTTTTTAATGGGAGAGACT BLK | Sigma |
| 14[207]12[208]BLK | AATTGAGAATTCTGTCCAGACGACTAAACCAA BLK | Sigma |
| 10[207]8[208]BLK | ATCCCAATGAGAATTAACTGAACAGTTACCAG BLK | Sigma |
| 6[207]4[208]BLK | TCACCGACGCACCGTAATCAGTAGCAGAACCG BLK | Sigma |
| 18[239]16[240]BLK | CCTGATTGCAATATATGTGAGTGATCAATAGT BLK | Sigma |
| 10[239]8[240]BLK | GCCAGTTAGAGGGTAATTGAGCGCTTTAAGAA BLK | Sigma |

**Supplementary table 10:** Staples with appropriate extensions for stem-loop configuration multiplexing

| Staple Name | Staple Sequence | Manufacturer |
|---|---|---|
| B2(1)22[47]20[48] R4x5 | CTCCAACGCAGTGAGACGGGCAACCAGCTGCA TTACACACACACACACA | Sigma |
| B6(1)14[47]12[48] R4x5 | AACAAGAGGGATAAAAATTTTTAGCATAAAGC TTACACACACACACACA | Sigma |
| B10(1)6[47]4[48] R4x5 | TACGTTAAAGTAATCTTGACAAGAACCGAACT TTACACACACACACACA | Sigma |
| F2(1)22[111]20[112] R4x5 | GCCCGAGAGTCCACGCTGGTTTGCAGCTAACT TTACACACACACACACA | Sigma |
| F6(1)14[111]12[112] R4x5 | GAGGGTAGGATTCAAAAGGGTGAGACATCCAA TTACACACACACACACA | Sigma |
| F10(1)6[111]4[112] R4x5 | ATTACCTTTGAATAAGGCTTGCCCAAATCCGC TTACACACACACACACA | Sigma |
| B2(2)22[175]20[176] R4x5 | ACCTTGCTTGGTCAGTTGGCAAAGAGCGGA TTACACACACACACACA | Sigma |
| B6(2)14[175]12[176] R4x5 | CATGTAATAGAATATAAAGTACCAAGCCGT TTACACACACACACACA | Sigma |
| B10(2)6[175]4[176] R4x5 | CAGCAAAAGGAAACGTCACCAATGAGCCGC TTACACACACACACACA | Sigma |
| F2(2)22[239]20[240] R4x5 | TTAACACCAGCACTAACAACTAATCGTTATTA TTACACACACACACACA | Sigma |
| F6(2)14[239]12[240] R4x5 | AGTATAAAGTTCAGCTAATGCAGATGTCTTTC TTACACACACACACACA | Sigma |
| F10(2)6[239]4[240] R4x5 | GAAATTATTGCCTTTAGCGTCAGACCGGAACC TTACACACACACACACA | Sigma |
| B2(1)22[47]20[48]R1x5 | CTCCAACGCAGTGAGACGGGCAACCAGCTGCA TTTCCTCCTCCTCCTCCTCCT | Sigma |
| B6(1)14[47]12[48]R1x5 | AACAAGAGGGATAAAAATTTTTAGCATAAAGC TTTCCTCCTCCTCCTCCTCCT | Sigma |
| B10(1)6[47]4[48]R1x5 | TACGTTAAAGTAATCTTGACAAGAACCGAACT TTTCCTCCTCCTCCTCCTCCT | Sigma |
| F2(1)22[111]20[112]R1x5 | GCCCGAGAGTCCACGCTGGTTTGCAGCTAACT TTTCCTCCTCCTCCTCCTCCT | Sigma |
| F6(1)14[111]12[112]R1x5 | GAGGGTAGGATTCAAAAGGGTGAGACATCCAA TTTCCTCCTCCTCCTCCTCCT | Sigma |
| F10(1)6[111]4[112]R1x5 | ATTACCTTTGAATAAGGCTTGCCCAAATCCGC TTTCCTCCTCCTCCTCCTCCT | Sigma |
| B2(2)22[175]20[176]R1x5 | ACCTTGCTTGGTCAGTTGGCAAAGAGCGGA TTTCCTCCTCCTCCTCCTCCT | Sigma |
| B6(2)14[175]12[176]R1x5 | CATGTAATAGAATATAAAGTACCAAGCCGT TTTCCTCCTCCTCCTCCTCCT | Sigma |
| B10(2)6[175]4[176]R1x5 | CAGCAAAAGGAAACGTCACCAATGAGCCGC TTTCCTCCTCCTCCTCCTCCT | Sigma |
| F2(2)22[239]20[240]R1x5 | TTAACACCAGCACTAACAACTAATCGTTATTA TTTCCTCCTCCTCCTCCTCCT | Sigma |

| | | |
|---|---|---|
| F6(2)14[239]12[240]R1x5 | AGTATAAAGTTCAGCTAATGCAGATGTCTTTC TTTCCTCCTCCTCCTCCTCCT | Sigma |
| F10(2)6[239]4[240]R1x5 | GAAATTATTGCCTTTAGCGTCAGACCGGAACC TTTCCTCCTCCTCCTCCTCCT | Sigma |

**Supplementary table 11:** Staples specific for stem-loop configuration

| Staple Name | Staple Sequence | Manufacturer |
|---|---|---|
| 11[160]12[144]_STEM_TAi_R5_R | CCAATAGCTCATCGTAGGAATCATGGCATCAA TT GAAGAAG TCGGCGAAAGCCGA | Sigma |
| 11[160]12[144]_STEM_GAP_R5_R | CCAATAGCTCATCGTAGGAATCATGGCATCAA TT GAAGAAG ACGGCGAAAGCCG | Sigma |

**Supplementary table 12:** Other staples used in the Photobleaching assay

| Staple Name | Staple Sequence | Manufacturer |
|---|---|---|
| B2(1)22[47]20[48] R5x5 | CTCCAACGCAGTGAGACGGGCAACCAGCTGCA TTCTTCTTCTTCTTCTTCTTC | Sigma |
| B6(1)14[47]12[48] R5x5 | AACAAGAGGGATAAAAATTTTTAGCATAAAGC TTCTTCTTCTTCTTCTTCTTC | Sigma |
| B10(1)6[47]4[48] R5x5 | TACGTTAAAGTAATCTTGACAAGAACCGAACT TTCTTCTTCTTCTTCTTCTTC | Sigma |
| F2(1)22[111]20[112] R5x5 | GCCCGAGAGTCCACGCTGGTTTGCAGCTAACT TTCTTCTTCTTCTTCTTCTTC | Sigma |
| F10(1)6[111]4[112] R5x5 | ATTACCTTTGAATAAGGCTTGCCCAAATCCGC TTCTTCTTCTTCTTCTTCTTC | Sigma |
| B2(2)22[175]20[176] R5x5 | ACCTTGCTTGGTCAGTTGGCAAAGAGCGGA TTCTTCTTCTTCTTCTTCTTC | Sigma |
| B6(2)14[175]12[176] R5x5 | CATGTAATAGAATATAAAGTACCAAGCCGT TTCTTCTTCTTCTTCTTCTTC | Sigma |
| B10(2)6[175]4[176] R5x5 | CAGCAAAAGGAAACGTCACCAATGAGCCGC TTCTTCTTCTTCTTCTTCTTC | Sigma |
| F2(2)22[239]20[240] R5x5 | TTAACACCAGCACTAACAACTAATCGTTATTA TTCTTCTTCTTCTTCTTCTTC | Sigma |
| F6(2)14[239]12[240] R5x5 | AGTATAAAGTTCAGCTAATGCAGATGTCTTTC TTCTTCTTCTTCTTCTTCTTC | Sigma |
| F10(2)6[239]4[240] R5x5 | GAAATTATTGCCTTTAGCGTCAGACCGGAACC TTCTTCTTCTTCTTCTTCTTC | Sigma |
| F6(1)14[111]12[112]S1 | GAGGGTAGGATTCAAAAGGGTGAGACATCCAA TTGTGATGTAGGTGGTAGAGGAA | Sigma |

**Supplementary table 13:** Staples for Stack-PAINT

| Staple Name | Staple Sequence | Manufacturer |
|---|---|---|
| 21[32]23[31]AA Stem | TTTTCACTCAAAGGGCGAAAAACCATCACC TT TCCTCCT TCGGCGAAAGCCGA | Sigma |
| 0[47]1[31]AA Stem | AGAAAGGAACAACTAAAGGAATTCAAAAAAA TT TCCTCCT TCGGCGAAAGCCGA | Sigma |
| 0[111]1[95]AA Stem | TAAATGAATTTTCTGTATGGGATTAATTTCTT TT TCCTCCT TCGGCGAAAGCCGA | Sigma |
| 21[120]23[127]AA Stem | CCCAGCAGGCGAAAAATCCCTTATAAATCAAGCCGGCG TT TCCTCCT TCGGCGAAAGCCGA | Sigma |
| 18[271]16[272]AA Stem | CTTTTACAAAATCGTCGCTATTAGCGATAG TT TCCTCCT TCGGCGAAAGCCGA | Sigma |
| 2[271]0[272]AA Stem | GTTTTAACTTAGTACCGCCACCCAGAGCCA TT TCCTCCT TCGGCGAAAGCCGA | Sigma |
| 21[32]23[31]AT Stem | TTTTCACTCAAAGGGCGAAAAACCATCACC TT TCCTCCT ACGGCGAAAGCCGT | Sigma |
| 0[47]1[31]AT Stem | AGAAAGGAACAACTAAAGGAATTCAAAAAAA TT TCCTCCT ACGGCGAAAGCCGT | Sigma |
| 0[111]1[95]AT Stem | TAAATGAATTTTCTGTATGGGATTAATTTCTT TT TCCTCCT ACGGCGAAAGCCGT | Sigma |
| 21[120]23[127]AT Stem | CCCAGCAGGCGAAAAATCCCTTATAAATCAAGCCGGCG TT TCCTCCT ACGGCGAAAGCCGT | Sigma |
| 18[271]16[272]AT Stem | CTTTTACAAAATCGTCGCTATTAGCGATAG TT TCCTCCT ACGGCGAAAGCCGT | Sigma |
| 2[271]0[272]AT Stem | GTTTTAACTTAGTACCGCCACCCAGAGCCA TT TCCTCCT ACGGCGAAAGCCGT | Sigma |
| 21[32]23[31]R1x3 | TTTTCACTCAAAGGGCGAAAAACCATCACC TT TCCTCCTCCTCCT | Sigma |
| 0[47]1[31]R1x3 | AGAAAGGAACAACTAAAGGAATTCAAAAAAA TT TCCTCCTCCTCCT | Sigma |
| 0[111]1[95]R1x3 | TAAATGAATTTTCTGTATGGGATTAATTTCTT TT TCCTCCTCCTCCT | Sigma |
| 21[120]23[127]R1x3 | CCCAGCAGGCGAAAAATCCCTTATAAATCAAGCCGGCG TT TCCTCCTCCTCCT | Sigma |
| 18[271]16[272]R1x3 | CTTTTACAAAATCGTCGCTATTAGCGATAG TT TCCTCCTCCTCCT | Sigma |

| | | |
|---|---|---|
| 2[271]0[272]R1x3 | GTTTTAACTTAGTACCGCCACCCAGAGCCA TT TCCTCCTCCTCCT | Sigma |

**Supplementary table 14:** Imaging Parameters

| Figure Number/ Data Sets | Laser Power ($W/cm^2$) | | Imager Concentrations (nM) | | Parameters (Exposure time, number of frames) | |
|---|---|---|---|---|---|---|
| | 561 nm | 640 nm | Cy3B | Atto647N | 561nm | 640nm |
| Figure 1d Left; Supp Figure-1a Left | 26 | 490 | 3 | 0.5 | 50 ms, 50000 | 100ms, 25000 |
| Figure 1d Right; Supp Figure-1a Right | 26 | 852 | 3 | 0.5 | 50 ms, 50000 | 100ms, 25000 |
| Figure 2 b, c; Supp Fig 2 | 26 | 490 | 3 | 0.5 | 50 ms, 50000 | 100ms, 25000 |
| Figure 3a(i); Supp Figure-3b (Replicate-1,3); Supp Figure-3c (Row 2); Supp Figure 4 (Replicate 1, 3) (Imagers: AGGAGGA-Cy3B, GAAGAAG-Atto647N) | 26 | 490 | 2.5 | 0.5 | 50 ms, 50000 | 100ms, 25000 |
| Figure 3a(iii); Supp Figure-3b (Replicate 1, 3); Supp Figure-3c (Row 2); Supp Figure 4 (Replicate 1, 3) (Imagers: TGTGTGT-Cy3B, AGGAGGA-Atto647N) | 26 | 490 | 2.5 | 0.5 | 50 ms, 50000 | 100ms, 25000 |
| Figure 3a(ii); Supp Figure-3b; Supp Figure-3c; Supp Figure 4 (Imagers: GAAGAAG-Cy3B; AGGAGGA-Atto647N) | 26 | 490 | 3 | 0.5 | 50 ms, 50000 | 100ms, 25000 |
| Figure 3a(iv); Supp Figure-3 b; Supp Figure-3c; Supp Figure 4 (Imagers: CTTCTTC-Cy3B; AGGAGGA-Atto647N) | 26 | 490 | 2.5 | 0.5 | 50 ms, 50000 | 100ms, 25000 |
| Supp Figure 3 b (Replicate 2); Supp Figure-3c (Row 1); Supp Figure 4 (Replicate 2) (Imagers: AGGAGGA-Cy3B, GAAGAAG-Atto647N) | 26 | 490 | 3 | 0.5 | 50 ms, 50000 | 100ms, 25000 |
| Supp Figure 3b (Replicate 2); Supp Figure-3c (Row 1); Supp Figure 4 (Replicate 2) (Imagers: TGTGTGT-Cy3B, AGGAGGA-Atto647N) | 26 | 490 | 5 | 0.5 | 50 ms, 50000 | 100ms, 25000 |
| Supp Figure 5b (Imagers: CTTCTTC-Cy3B; AGGAGGA-Cy3B, TGTGTGT-Cy3B) | 26 | 490 | 2.5, 0.5, 0.5 respectively | | 50 ms, 50000; 100ms, 10000 | |
| Supp Figure 5c (Imagers: CTTCTTC-Cy5; AGGAGGA-Cy3B) | 313 | 57 | 0.5 | 1.5 (Cy5) | 100 ms, 25000 | 50 ms, 50000 |
| Supp Figure 6 b, c | 26 | | | | 200 ms, 15000 | |
| Figure 4, Supp Figure 10, 11 (Imagers: AGGAGGA-Cy3B, GAAGAAG-Atto647N) | | | 0.5 | 0.5 | 50ms, 72000 frames | 100ms, 25000 frames |
